# Adaptive tuning of human learning and choice variability to unexpected uncertainty

**DOI:** 10.1101/2022.12.16.520751

**Authors:** Junseok K. Lee, Marion Rouault, Valentin Wyart

## Abstract

Human value-based decisions are strikingly variable under uncertainty. This variability is known to arise from two distinct sources: variable choices aimed at exploring available options, and imprecise learning of option values due to limited cognitive resources. However, whether these two sources of decision variability are tuned to their specific costs and benefits remains unclear. To address this question, we compared the effects of expected and unexpected uncertainty on decision-making in the same reinforcement learning task. Across two large behavioral datasets, we found that humans choose more variably between options but simultaneously learn less imprecisely their values in response to unexpected uncertainty. Using simulations of learning agents, we demonstrate that these opposite adjustments reflect adaptive tuning of exploration and learning precision to the structure of uncertainty. Together, these findings indicate that humans regulate not only how much they explore uncertain options, but also how precisely they learn the values of these options.

**Teaser:** Humans regulate not only how much they explore uncertain options, but also how precisely they learn their values.

## Introduction

Human decisions exhibit a pervasive variability under uncertainty (*1*). In the context of value-based decisions, the source of this variability has classically been assigned to exploration (*2*–*5*) – i.e., purposeful bias and variance in choice policy aimed at reducing uncertainty about the values of choice options. Beside this well-described source of decision variability, recent work has shown that the computations used to learn option values from obtained rewards suffer from imprecisions due to limited cognitive resources (*6*–*9*). Learning imprecisions result in decision variability which can be mistaken for exploration, but these two sources of decision variability are dissimilar in nature. Exploration drives decision variability through the probabilistic selection of options which do not maximize expected value, whereas learning imprecisions reflect random noise in the reinforcement learning process that updates option values. In practice, these differences allow decomposing human decision variability into two separate sources through detailed computational modeling of human behavior (*6*).

Both exploration and imprecise computations entail significant reward costs. First, by selecting an option that does not maximize expected value, exploration temporarily foregoes the exploitation of the best available option (*10*). Second, relying on imprecise computations means that the option with the highest subjective value is less likely to be the objectively best option available. Despite these similar reward costs, the two sources of decision variability have very different cognitive benefits. Exploration reduces uncertainty about option values, whereas imprecise learning reduces demands in terms of cognitive and neural resources (*11*–*13*). It is well known that humans arbitrate the ‘explore-exploit’ trade-off under uncertainty in terms of its costs and benefits (*3, 4*). However, these findings have been obtained using reinforcement learning models which assign all decision variability to exploration. Whether humans simultaneously regulate learning imprecisions in terms of their specific costs and benefits remains unknown.

Importantly, the costs and benefits of exploration and learning imprecisions depend on the dominant form of uncertainty in the environment: expected vs. unexpected uncertainty (*14, 15*). Expected uncertainty refers to random stochasticity of rewards associated with a choice option around a constant mean value. By contrast, unexpected uncertainty refers to changes in the mean value of rewards associated with a choice option. Under expected uncertainty (i.e., reward stochasticity), individual rewards become less informative about their mean value as learning progresses, and agents can therefore tolerate low learning rates and little exploration. By contrast, under unexpected uncertainty (i.e., reward volatility), individual rewards are highly informative about changes in their mean value, and agents should therefore maintain high learning rates and frequent exploration of unobserved choice alternatives. There is ample experimental evidence that humans adjust their learning rates and exploration at short timescales depending on the dominant form of uncertainty in their environment (*16*–*19*). However, whether this regulation of learning rates and exploration is accompanied by a modulation of learning imprecisions remains unknown.

Here, we developed a novel experimental framework in which the effects of expected and unexpected uncertainty on decision variability can be compared in the context of the same reinforcement learning task (Figure 1AB; see *Materials and Methods*). We decomposed the decision variability of two large samples of human participants (a ‘discovery’ dataset and a ‘replication’ dataset; see *Materials and Methods*) into exploration and learning imprecisions, by fitting a noisy reinforcement learning model to their behavior. We found that participants choose more variably between options but learn more precisely their values in response to unexpected uncertainty. By studying individual differences in these opposite adjustments, we show that humans regulate the variability of their decisions based on not one, but two cost-benefit trade-offs.

**Figure 1.**
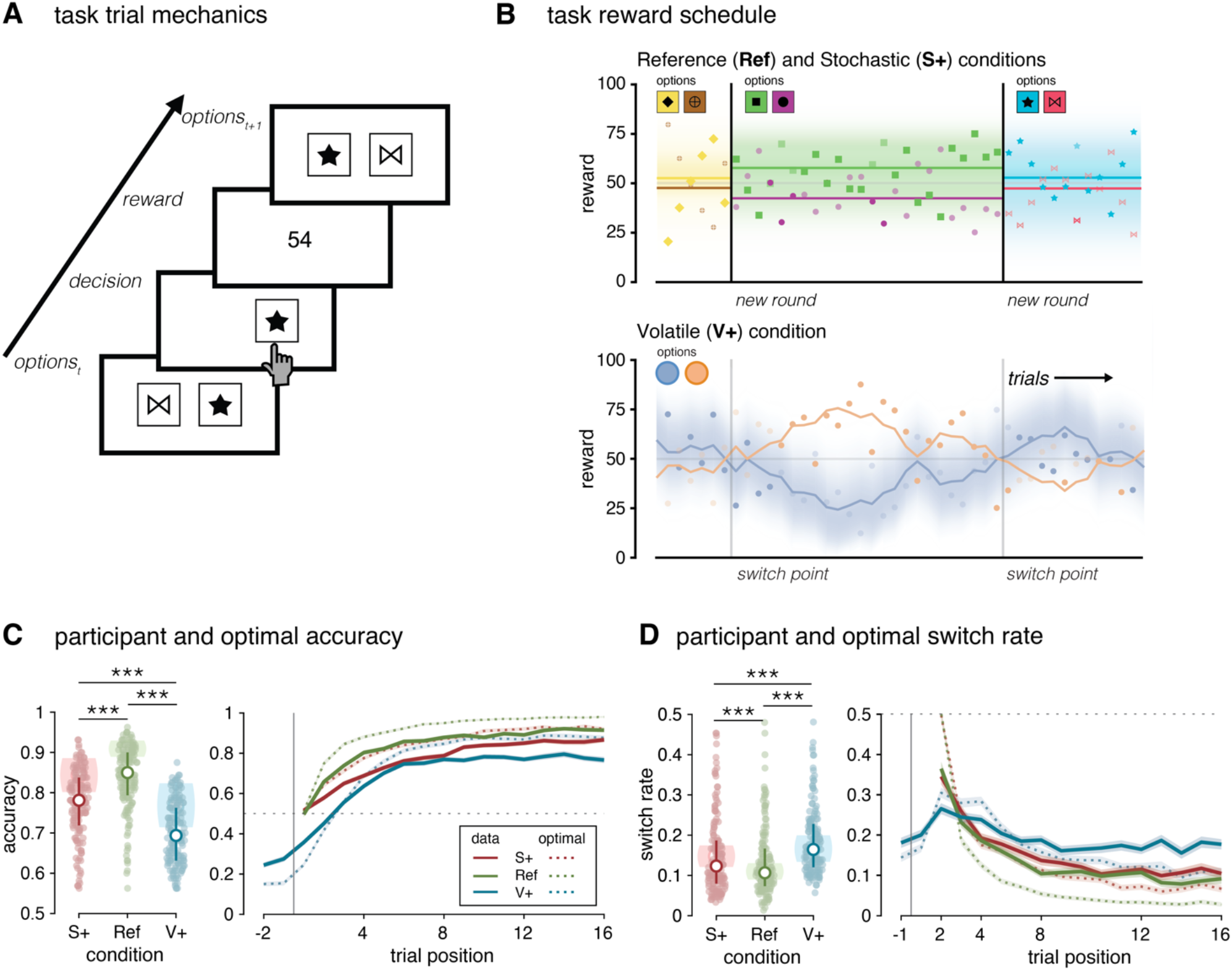
Experimental task properties and behavioral results. (**A**) Trial modalities. Two options are shown to the participant. Once a choice is made, the reward associated with the option on that trial is displayed. (**B**) Reward schedule of an option during a block. Colored shapes indicate the reward given for an option. The set of shape options are shown for each block. Bold colored shapes indicate the option chosen on a trial. Vertical lines demarcate the beginning and the end of a block in the Reference (Ref) and the Stochastic (S+) conditions and a reversal in the Volatile (V+) condition. Upper: Bold red horizontal lines show the generative mean for the option corresponding to the correct shape in the Reference and Stochastic conditions. Lower: Bold lines indicate the generative reward mean for the best option drifting throughout the course of a block in the Volatile condition. Shaded areas around lines represent the values of the probability density function from which a reward (shape) was drawn. (**C**) Left: Average accuracy within each condition. Colored dots represent individual participants’ mean accuracy. White dots indicate the median accuracy. Error bars represent the 1st and 3rd quartiles. Shaded areas indicate 1st and 3rd quartiles of the optimal model’s accuracy. Right: Accuracy over time within each condition. The vertical line represents the start of a new block or a reversal. Solid lines indicate the mean accuracy across participants. Dotted lines indicate the mean accuracy of the optimal model. Shaded areas correspond to the SEM. (**D**) Left: Average switching rate within each condition. The vertical line represents the start of a new block or a reversal. Colored dots represent individual participants’ overall switch rate. White dots indicate the median switch rate. Error bars represent the 1st and 3rd quartiles. Shaded areas indicate 1st and 3rd quartiles of the optimal model’s switch rate. Right: Proportion of switches over time within each condition. Solid lines indicate the mean switch rate across participants. Dotted lines indicate the mean accuracy of the optimal model. Shaded areas correspond to the SEM.

## Results

### Two-armed bandit task design and performance

Adult participants (*N* = 200 per dataset; see *Materials and Methods*) played a two-armed bandit task in three conditions (Figure 1B). The ‘reference’ (Ref) condition consists of short rounds of trials using two choice options associated with reward distributions of fixed means and variances. The ‘stochastic’ (S+) condition differs from the reference condition in that reward distributions have larger variances – i.e., increased expected uncertainty. The ‘volatile’ (V+) condition differs from the reference condition in that it consists of long rounds of trials with reward distributions of changing means – i.e., unexpected uncertainty. Participants were instructed of the structure of uncertainty and relative difficulty of each condition (see *Materials and Methods*). We describe below the analysis of the first, ‘discovery’ dataset of adult participants (*N* = 154 after exclusion; see *Materials and Methods*) and point toward the replication of observed effects in the second, ‘replication’ dataset (*N* = 142 after exclusion; see *Materials and Methods*).

As expected, participants were more accurate at choosing the option associated with the highest reward mean in the reference condition than in the other two conditions with increased uncertainty (Figure 1C; signed-rank test, S+ minus Ref: *z*= -7.1, *p* < 0.001, V+ minus Ref: *z*= -10.2, *p* < 0.001). Participants also switched more often between choice options in the stochastic and volatile conditions than in the reference condition (Figure 1D; S+ minus Ref: *z*= 3.9, *p* < 0.001, V+ minus Ref: *z*= 8.0, *p* < 0.001). Comparing the two conditions with increased uncertainty showed that participants switched more often between choice options in the volatile condition (V+ minus S+: *z*= 5.3, *p* < 0.001). These differences between conditions were replicated in the second dataset (Supplementary Figure 1).

### Computational model specification

We first sought to compare human decisions to those of an optimal learning agent in the same task conditions. For this purpose, we derived a Kalman filter whose parameters were set to the generative values of each task condition (see *Materials and Methods*). The optimal learning agent selected on each trial of each round the option associated with the highest estimated value (reward mean). Simulating the behavior of the optimal learning agent confirmed that human decisions were substantially less accurate and more variable than those of the optimal learning agent in all three conditions (Figure 1CD). These differences between human and optimal decisions were replicated in the second dataset (Supplementary Figure 1).

To capture the suboptimal variability of human decisions, we altered the optimal learning agent with four free parameters (Figure 2AB; see *Materials and Methods*). First, a learning rate *α* controls how much estimated option values are updated following each obtained reward. Unlike reinforcement learning (RL) models, the learning rate *α* of a Kalman filter reflects the dominant form of uncertainty assumed by the learning agent, and the effective magnitude of updates varies from trial to trial as a function of uncertainty regarding the current value of the chosen option (see *Supplementary Text*). Second, a decay rate *δ* controls the rate at which the value of the unchosen option regresses toward its prior value, reflecting a decay of unchosen option values in working memory. Third, a learning noise term triggers imprecise updates of estimated option values, controlled by a scaling factor *ζ*. As in recent work (*6*), we hypothesized that learning imprecisions follow Weber’s law, and scale with the magnitude of associated reward prediction errors. And fourth, a choice temperature *τ* generates exploration through a ‘softmax’ choice policy (*3, 4*). Before analyzing fits of this suboptimal learning agent to human decisions, we performed model and parameter recovery analyses (*20, 21*) to validate that our fitting procedure was capable of identifying the source(s) of decision variability from choice behavior (Supplementary Figure 2; see *Materials and Methods*). We also verified that the main findings are robust to the use of a reinforcement learning (RL) model instead of a Kalman filter to fit human decisions (see *Supplementary Text*).

**Figure 2.**
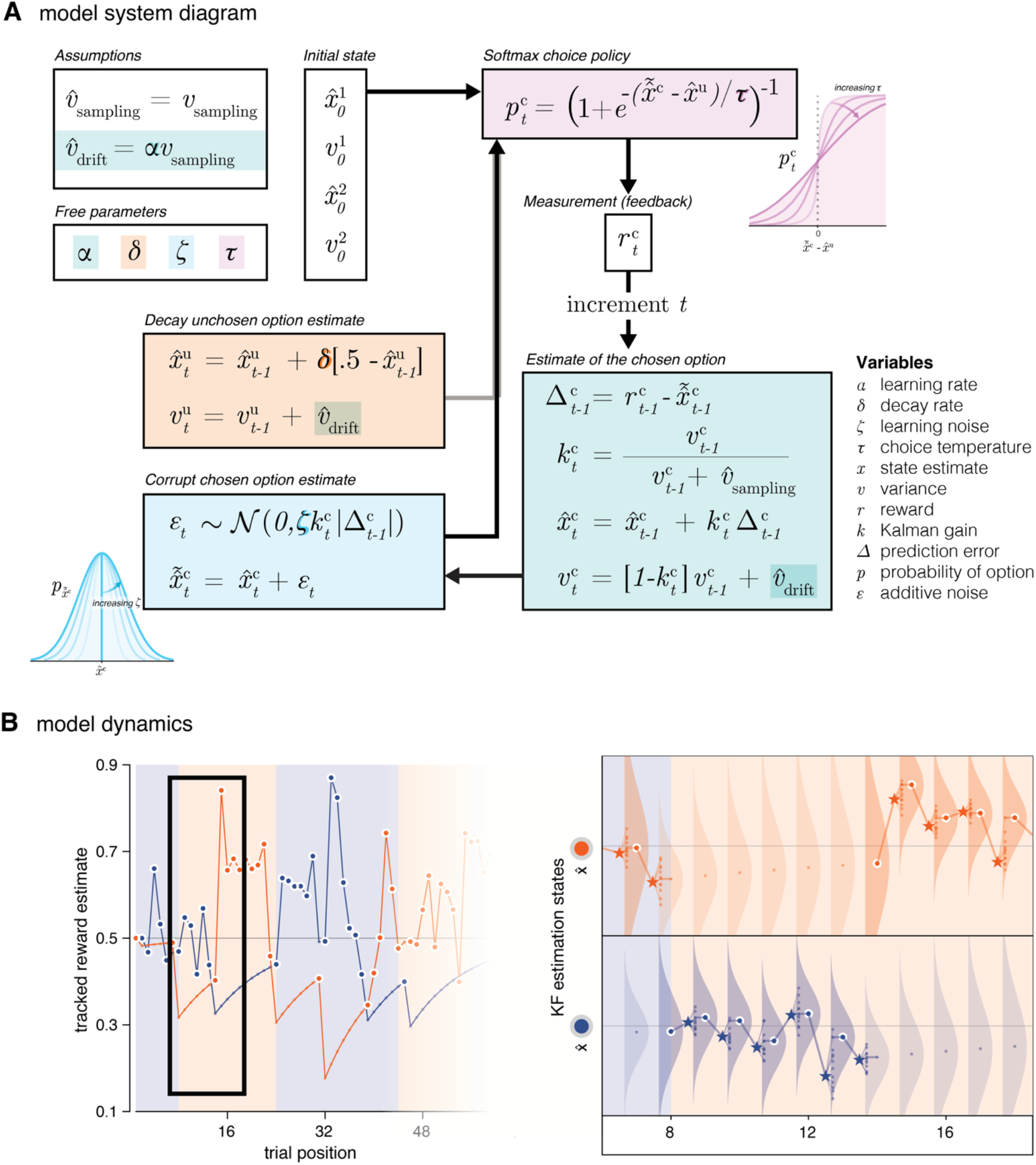
Details of the noisy Kalman Filter model. (**A**) System diagram of the noisy KF model. The model tracks the means *x*_*t*_ and variances *v*_*t*_ of option values based on the sampling variance *v*_*s*ampling_ and drift variance *v*_*d*rif*t*_ of observed rewards r_*t*_. The model includes four free parameters, which control the update of option values (*α*, green box), the corruption of option values by learning noise (*ζ*, blue box), the decay of unchosen option values to their overall mean (*δ*, orange box), and the softmax choice policy (*τ*, purple box). Insets illustrate the effects of the learning noise and choice temperature parameters on the distributions of option value and decision probability, respectively. (**B**) Dynamics of the model within the task context (simulated on the V+ condition). Left: The trajectory of the estimated reward for each option over time. Chosen options are highlighted with a white border. Right: Detailed view of the model dynamics. Dots correspond to the tracked values in the left panel. Stars correspond to the reward feedback after choice. Smaller dots following the feedback correspond to potential learned values corrupted by learning noise. Distributions around dots correspond to the estimation uncertainty. Note: this estimation uncertainty increases for unchosen options as the tracked value itself decays over time.

### Model parameter fits in stochastic and volatile conditions

We fitted the parameters of the suboptimal learning agent to each participant’s behavior in each condition using approximate Bayesian inference (see *Materials and Methods*). We performed Bayesian model selection to compare the suboptimal learning agent including the two sources of decision variability (learning imprecisions and exploration) with variants including a single source (Supplementary Figure 3). We found that both sources of decision variability are needed to fit participants’ behavior (exceedance *p* > 0.999 in each condition). In line with recent work, we validated that learning imprecisions scale with the magnitude of updates following each outcome (comparison between flat and update-scaled imprecisions, exceedance *p* > 0.999 in each condition). We found also evidence for a significant memory decay of unchosen option values in all three conditions (comparison between *δ* > 0 and *δ* = 0, exceedance *p* > 0.999 in each condition).

As expected, participants had higher learning rates in the volatile condition than in the other two conditions (Figure 3A; V+ against Ref: *z*= 10.2, *p* < 0.001, V+ against S+: *z*= 9.8, *p* < 0.001). Regarding decision variability, participants had lower learning noise in the volatile condition (V+ against Ref: *z*= -2.8, *p* = 0.006, V+ against S+: *z*= -4.4, *p* < 0.001), as well as higher choice temperature (V+ against Ref: *z*= 4.4, *p* < 0.001, V+ against S+: *z*= 4.4, *p* < 0.001). Importantly, we found no statistically significant difference between the reference and stochastic conditions in terms of any model parameter (learning rate: *p* = 0.437 decay rate: *p* = 0.479, learning noise: *p* = 0.100, choice temperature: *p* = 0.995). Unlike its optimal counterpart, this suboptimal learning agent was able to capture participants’ behavior in terms of accuracy and switch rate, as well as the trajectories of these quantities over the course of each round (Figure 3BC). These effects were replicated in the second dataset (Supplementary Figure 1).

**Figure 3.**
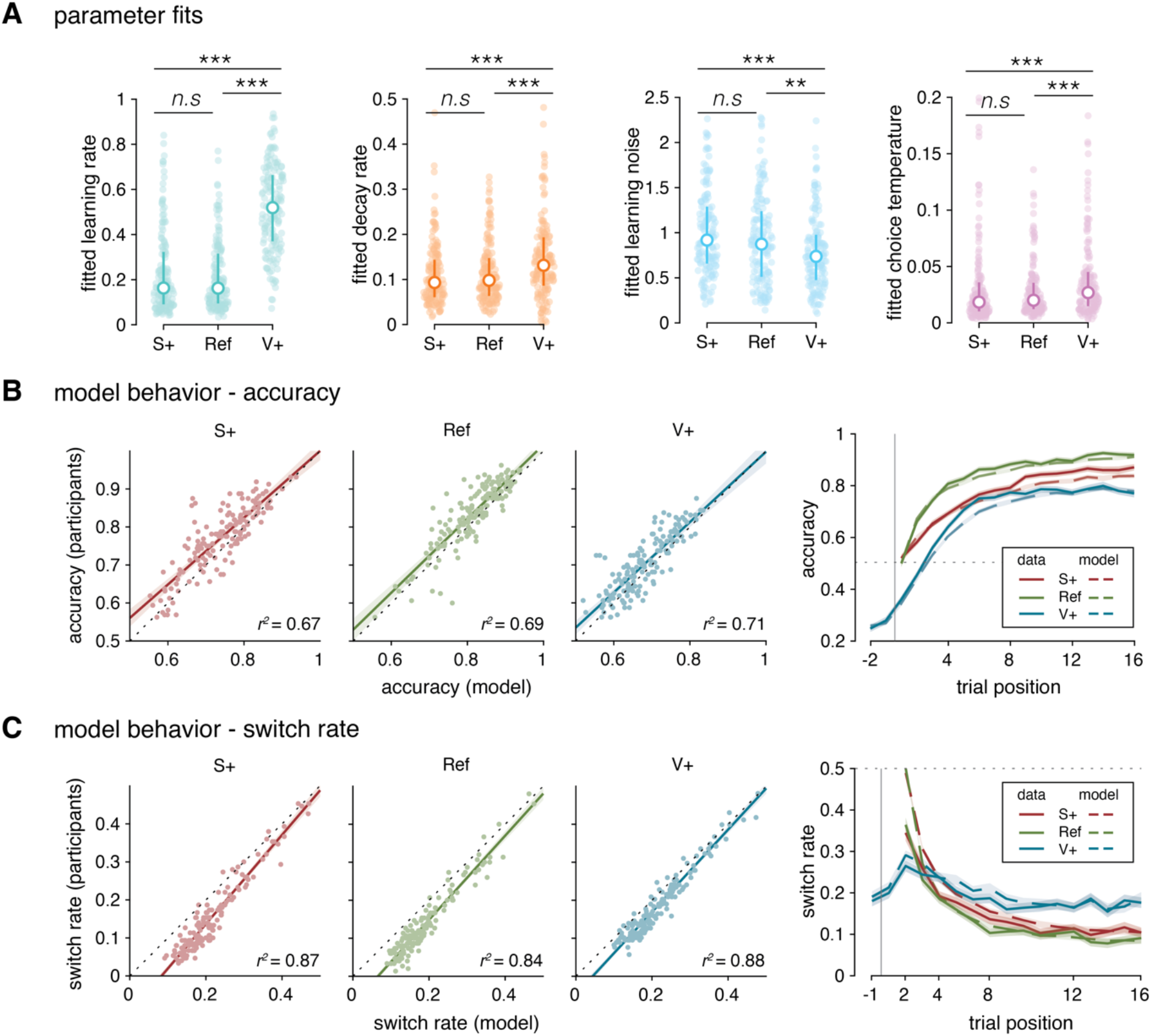
Summary of model fits to participant data. (**A**) Colored dots represent individual participants’ fits for each parameter. White dots indicate the median fitted parameter value. Error bars represent the 1st and 3rd quartiles. (**B**) Left: Scatter plots of the average accuracy of participants to their best fitting models in each condition. Right: Accuracy over time within each condition. Vertical line represents the start of a new block or a reversal. Solid lines indicate the mean accuracy across participants. Dashed lines indicate the mean accuracy of the best fitting model. (**C**) Left: Scatter plots of the switch rate of participants to their best fitting models in each condition. Right: Proportion of switches over time within each condition. Vertical line represents the start of a new block or a reversal. Solid lines indicate the mean switch rate across participants. Dotted lines indicate the mean accuracy of the best fitting model. Shaded areas correspond to the SEM.

Learning noise and choice temperature showed large individual differences in each condition. We took advantage of these differences to study how each parameter impacts decision variability. We correlated each parameter with decision accuracy and switch rate across participants (Figure 4A). Participants’ accuracy showed significant negative correlations with learning imprecisions and exploration (rank correlation, learning noise: *ρ* = -0.18, *p* < 0.001, choice temperature: *ρ* = -0.38, *p* < 0.001). By contrast, participants’ switch rates showed a negative relation with learning noise (*ρ* = -0.14, *p* = 0.002), but a strong positive relation with choice temperature (*ρ* = 0.71, *p* < 0.001). This means that unlike exploration, learning imprecision did not make participants switch more between the two choice options. Median splits of participants’ choice behavior as a function of learning noise (Figure 4B) or choice temperature (Figure 4C) confirmed these effects, which were replicated in the second dataset (Supplementary Figure 3).

**Figure 4.**
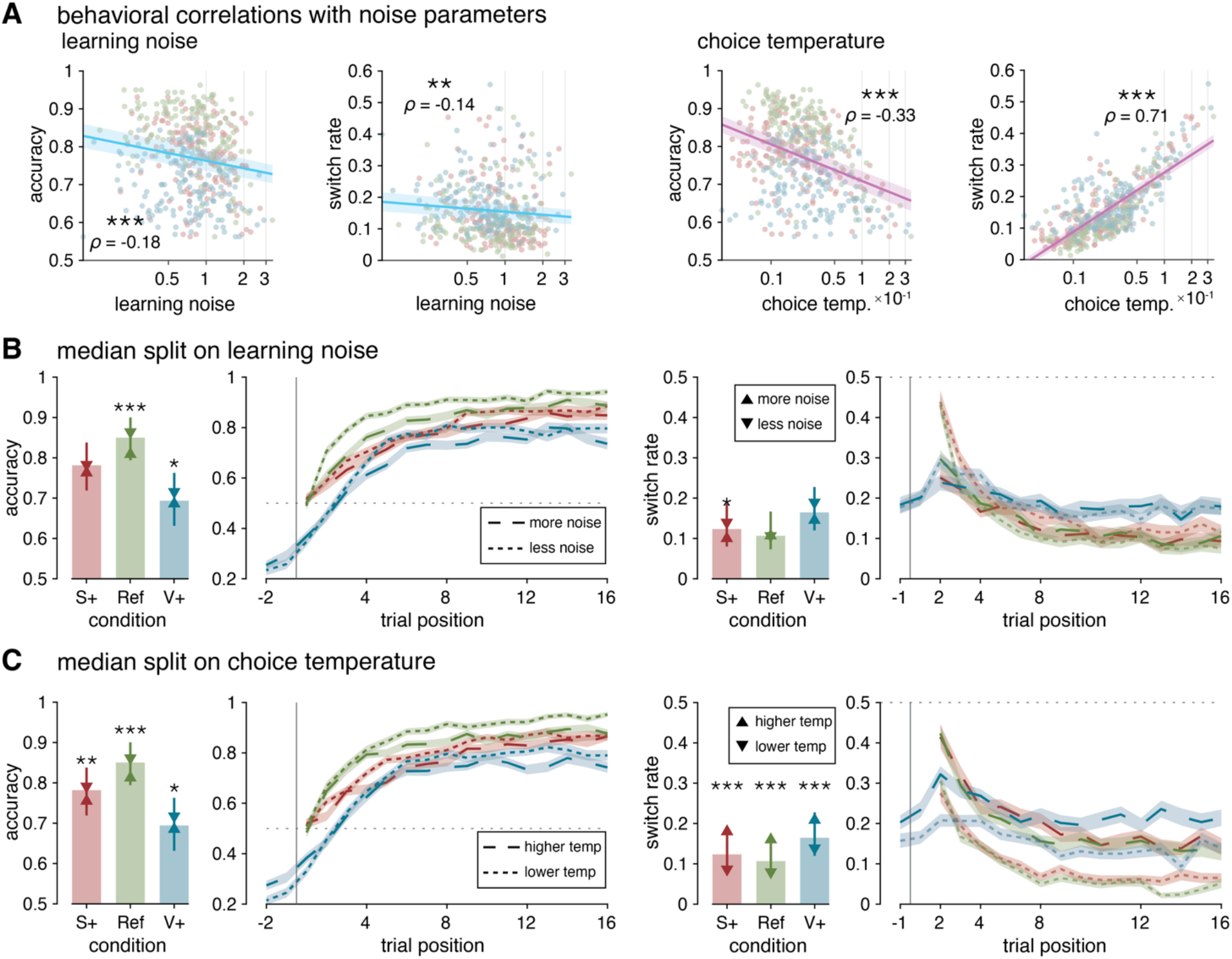
Behavioral expressions of learning and decision noise. (**A**) Correlations of behavioral measures with learning noise and choice temperature. Learning noise and choice temperature axes are spaced with log-scaling. Shaded areas are 95% CI. (**B**) Participant accuracy and switch rates with respect to learning noise (median split across participants’ best-fitting learning noise in each of the three conditions separately). Statistical significance determined from rank-sum tests on median split participants. (**C**) Participant accuracy and switch rates with respect to choice temperature (median split across participants’ best-fitting choice temperature in each of the three conditions separately). Error bars represent interquartile ranges. Upward triangles and solid lines signify the mean of the upper values of the measure on the median split of the parameter. Downward triangles and dashed lines signify the mean of the measure on the lower values of the median split of the parameter.

### Model parameter covariations across participants

We studied how the large individual differences in decision variability are shared between model parameters and across conditions. For this purpose, we computed the correlation matrix of model parameters across participants (12 parameters = 4 parameters per condition; see *Materials and Methods*). This matrix revealed significant covariations between model parameters and conditions (Figure 5A). First, learning rate, learning noise and choice temperature all showed significant within-parameter correlations between the reference and the stochastic or volatile conditions (Figure 5C). Furthermore, learning rate correlated negatively with learning noise (Figure 5D; rank correlation, *ρ* = -0.36, *p* < 0.001), but positively with choice temperature (*ρ* = 0.43, *p* < 0.001). We verified that these between-parameter correlations remained significant within each condition, and performed additional recovery analyses to validate that neither reflects spurious correlations arising from the fitting procedure (Figure 5B; see *Materials and Methods*). These correlations were also replicated in the second dataset (Supplementary Figures 4 and 5).

**Figure 5.**
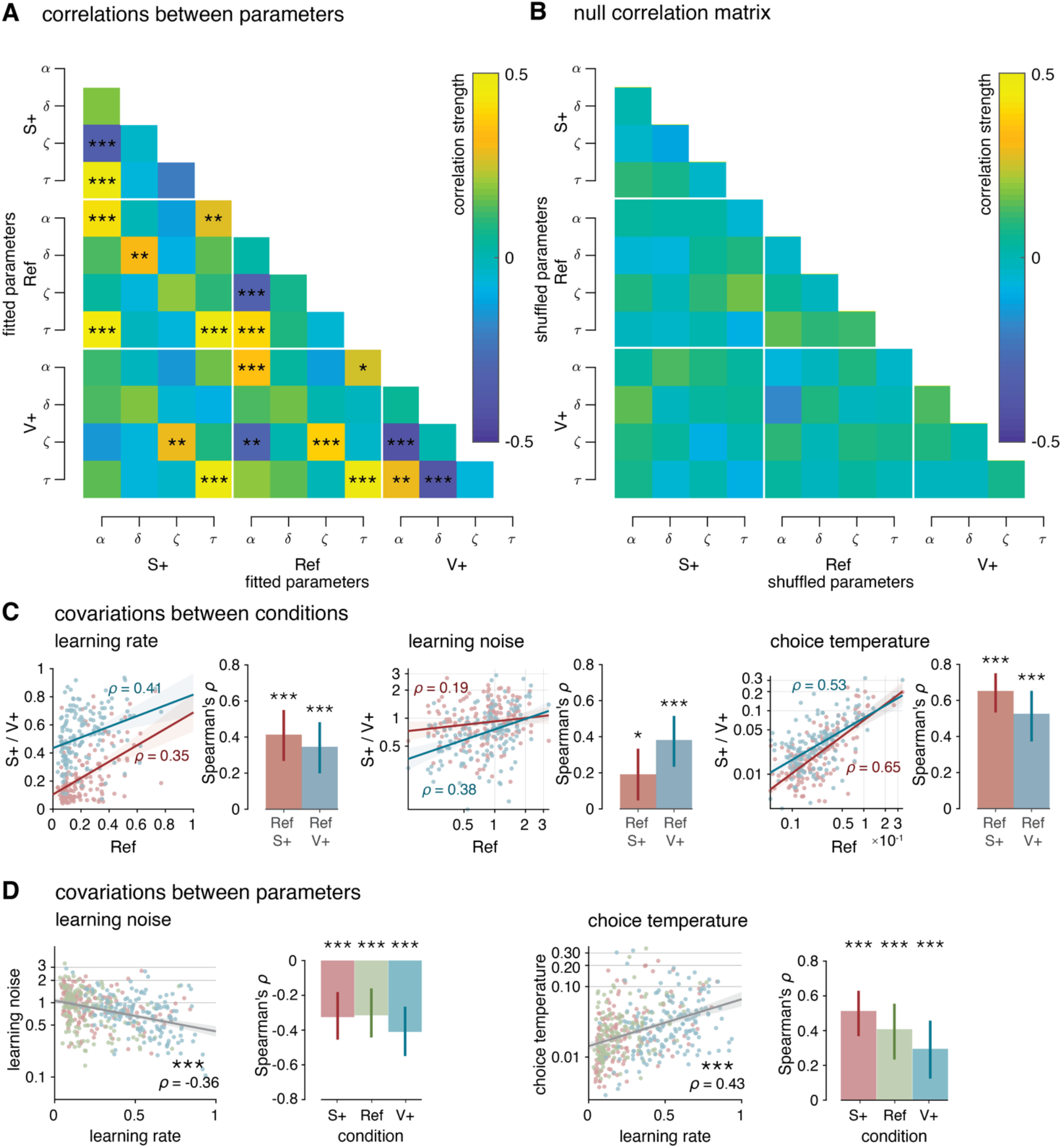
Model parameter covariations across participants. (**A**) Correlation matrix of participants’ fitted parameters. (**B**) Null correlation matrix (correlation structure destroyed) using shuffled parameters for simulations, then fitted using the same procedure. All p-values in correlation matrices corrected for false discovery rates (*α* = 0.05). (**C**) Covariations of the learning rate, learning noise, and choice temperature between the reference (R) to the stochastic (S+) and the volatile (V+) conditions. Lines in scatter plots indicate the best-fitting regression line (blue: V+ to R; red: S+ to R). Shaded areas are 95% CI. Error bars on bar plots indicate 95% bootstrapped CI on the Spearman’s *ρ* value. (**D**) Correlations of learning noise and choice temperature with learning rate. Regression lines fitted on aggregate data pooling all conditions. Shaded areas are 95% CI. Error bars on bar plots indicate 95% bootstrapped CI on the Spearman’s *ρ* value.

The positive association of learning noise and learning rate as well as the negative association of choice temperature and learning rate across participants mirror their within-participant adjustments across task conditions. Indeed, the decrease in learning noise and the increase in choice temperature are associated with higher learning rate in the volatile condition. But are these simultaneous adjustments of decision variability distinct or tied to each other? To answer this question, we first measured the partial correlation between learning noise and choice temperature once the relation of these two quantities with learning rate are partialled out. This analysis indicated no direct relation between the two quantities (partial rank correlation, *ρ* = 0.03, *p* = 0.584). We then measured the relation between adjustments of each source of variability with adjustments of learning rate between conditions. We found that the decrease in learning noise in the volatile condition (V+ minus Ref) correlates with the increase in learning rate (*ρ* = -0.22, *p* = 0.007), but not with the increase in choice temperature (*ρ* = -0.03, *p* = 0.739). Together, these results indicate that the opposite adjustments of learning noise and choice temperature to volatility are independent of each other.

### Principal component analysis of model parameters

To provide additional support for independent adjustments of learning imprecision and exploration to volatility, we performed a principal component analysis (PCA) of model parameters (see *Materials and Methods*). By construction, principal components (PCs) are orthogonal to each other and reflect uncorrelated sources of variability in model parameters. We therefore asked whether learning noise and choice temperature project onto distinct PCs. This analysis revealed two PCs (PC1 and PC2) that capture covariance in model parameters better than chance (Figure 6A).

**Figure 6.**
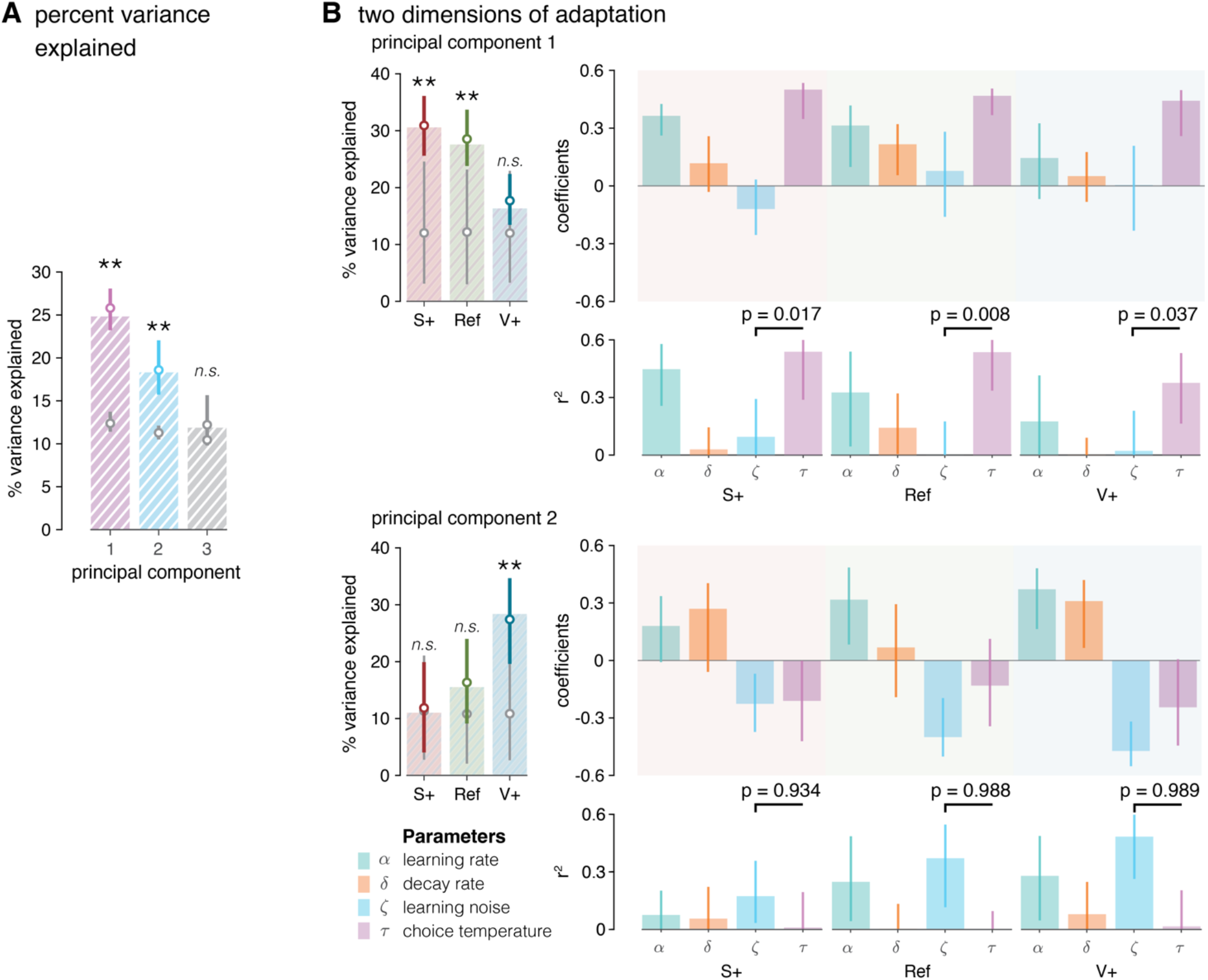
Principal component analysis of model parameters. (**A**) Percent variance explained up to the first non-contributive PC. Colored dots correspond to the median value of the percent variance explained from the bootstrap procedure. Gray dots are median values of the percent variance explained from PCs of shuffled data. Statistical significance calculated from one-tailed bootstrap significance tests. (**B**) Left: Percent variance explained by the first two PCs within each condition. The first PC, dominated by the variation of choice temperature, explains best the parameter adaptations in the non-volatile conditions. The second PC, dominated by the variation of learning noise, explains best the parameter adaptations in the volatile condition. Upper right: Ingredients and coefficients of the first two principal components. Lower right: Coefficient of determination of each parameter for the PC. All error bars indicate 95% bootstrapped CI.

PC1 reflected the positive correlation between learning rate and choice temperature, and explained more variance in model parameters in the reference (Ref) and stochastic (S+) conditions (Figure 6B, top). Importantly, PC1 explained significantly more variance in choice temperature than learning noise in all three conditions (Ref: bootstrapped *p* = 0.992, S+: bootstrapped *p* = 0.983, V+: bootstrapped *p* = 0.963). Behaviorally, high PC1 scores were associated with lower accuracy and higher switch rates (see *Supplementary Text*). By contrast, PC2 (Figure 6B, bottom) reflected the negative correlation between learning rate and learning noise, and explained more variance in model parameters in the volatile (V+) condition. Importantly, PC2 explained significantly more variance in learning noise than choice temperature in all conditions (Ref: bootstrapped *p* = 0.988, S+: bootstrapped *p* = 0.934, V+: bootstrapped *p* = 0.989). Behaviorally, PC2 showed a milder relation to accuracy and no significant relation to switch rate (see *Supplementary Text*). These different effects were replicated in the second dataset (Supplementary Figure 6). The covariance structure of model parameters extracted using PCA confirms that the simultaneous adjustments of learning noise and choice temperature to volatility are largely independent of each other.

### Adaptive regulation of learning and choice variability to uncertainty

Participants choose more variably but learn less imprecisely in the volatile condition as compared to the other two conditions. But do these opposite adjustments of decision variability reflect the changing costs of learning imprecisions and exploration across conditions? To address this question, we performed theoretical simulations of noisy reinforcement learning agents to measure the reward costs of learning imprecisions and exploration in each condition.

We computed the marginal reward costs associated with each source of decision variability through simulations of the suboptimal learning agent. We selectively varied the associated model parameter (either the learning noise *ζ* or the choice temperature *τ*) while holding all other model parameters constant and set to their best-fitting values (see *Materials and Methods*). We measured marginal reward cost *C*_*x*_ as the fraction loss of reward excess (i.e., the difference between obtained and foregone rewards) compared to a learning agent without the corresponding source of variability *x* (*x* = *ζ* or *τ*) tested in the same condition (Ref, S+ or V+). This ‘relative’ definition of reward cost *C* was chosen to measure how much each source of variability impacts the reward that could have been obtained in each condition (from *C*_*x*_ = 0 for a cost-free source of variability to *C*_*x*_ = 100% for a purely random agent).

Marginal reward costs *C*_>_ and *C*_*τ*_ increased monotonically with learning noise and choice temperature in all conditions (Figure 7A). However, the same amount of learning noise *ζ* was associated with larger reward costs *C*_>_ in the volatile condition than in the other two conditions. By contrast, the same choice temperature *τ* was associated with smaller reward costs *C*_*τ*_ in the volatile condition. This means that learning noise is effectively more costly in the volatile condition, whereas choice temperature is effectively less costly in the volatile condition.

**Figure 7.**
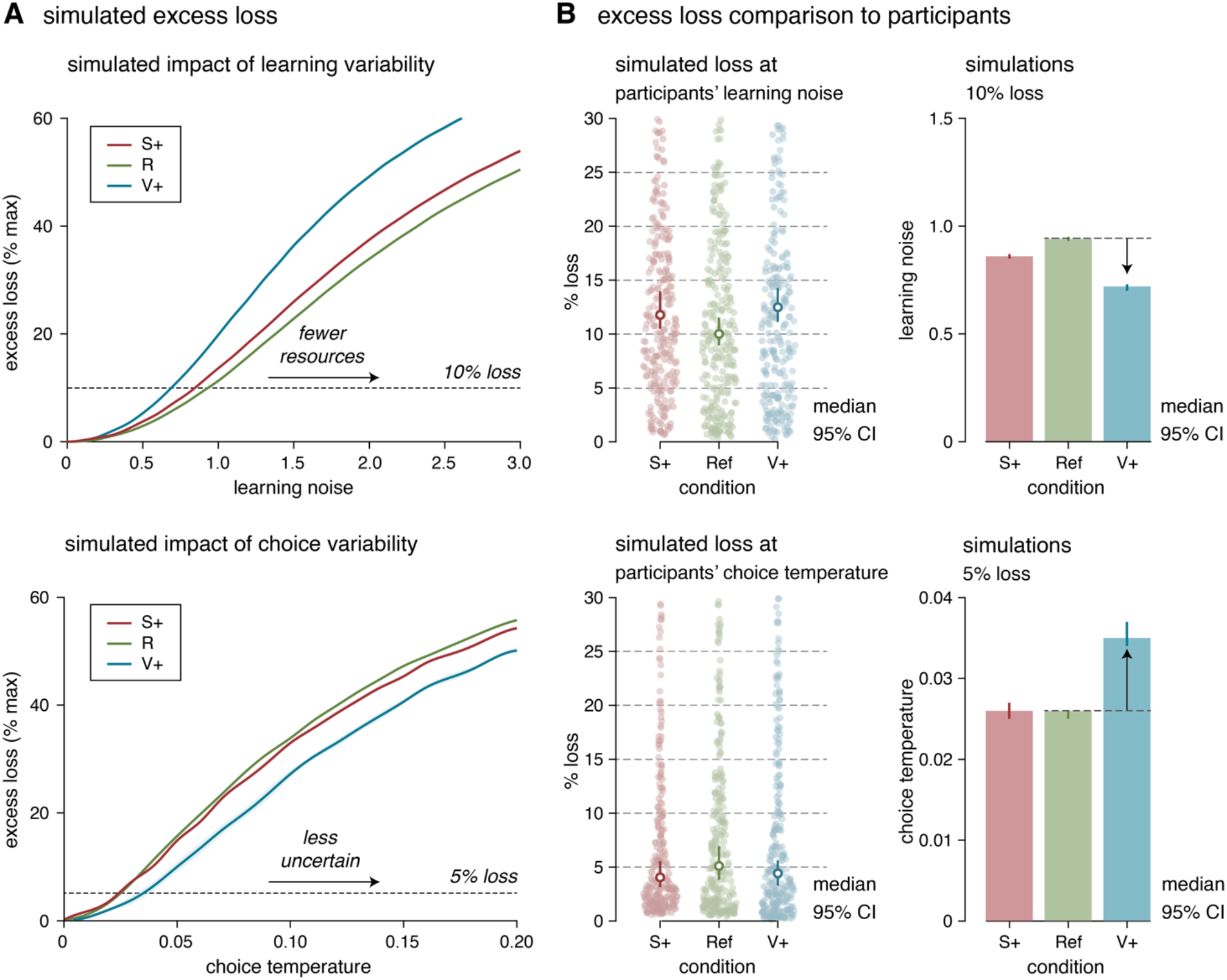
Adaptive adjustment of model parameters to uncertainty. (**A**) Marginal reward costs (reward loss, expressed as % of maximum obtainable reward rate) associated with each source of decision variability through simulations of the suboptimal learning agent, by varying selectively the associated model parameter (either the learning noise *ζ* or the choice temperature *τ*) while holding all other model parameters constant and set to their best-fitting values. Top: the same amount of learning noise is associated with larger marginal reward cost in the volatile (V+) condition. Bottom: the same choice temperature is associated with smaller marginal reward cost in the volatile (V+) condition. Inset: marginal reward costs associated with observed amounts of learning noise (top) and choice temperatures (bottom) in each of the three conditions. (**B**) Simulated vs. observed adjustments. Top: amounts of learning noise associated with the same marginal reward cost (here, 10% reward loss) in the three conditions. Obtained values (left) match observed learning noise estimates in participants’ data (right). Bottom: choice temperatures associated with the same marginal reward cost (here, 5% reward loss) in the three conditions. Obtained values (left) match observed choice temperature estimates in participants’ data (right).

Based on these theoretical simulations, we estimated the marginal reward costs incurred by learning noise (*C*_>_) and choice temperature (*C*_*τ*_) for each participant and each condition (Figure 7B; *N* = 296 across the two datasets). We found that *C*_>_ did not differ between the S+ and V+ conditions (S+: 11.8% [10.5 13.9]; V+: 12.5% [10.9 14.3], median [bootstrapped 95% CI]; *z*= 1.0, *p* = 0.307). Like *C*_>_, *C*_*τ*_ did not differ between these two conditions (S+: 4.1% [3.2 5.6]; V+: 4.4% [3.4 5.6]; z = 1.3, *p* = 0.190). In other words, the decrease in learning noise and increase in choice temperature in the volatile condition compensate for the larger reward costs of learning noise and the smaller reward costs of choice temperature in this condition. Importantly, expressing reward costs in terms of ‘joint’ reward costs of the two sources of variability *C*_>,*τ*_ (compared to a learning agent without any variability) also results in similar costs across conditions (Supplementary Figure 7C).

Conversely, the amounts of learning noise *ζ* and choice temperature *τ* associated with fixed marginal reward costs across conditions show a decrease in *ζ* and an increase in *τ* in the volatile condition – as in participants (Figure 7B). In additional theoretical analyses (see *Supplementary Text*), we considered alternative definitions of reward costs and the possibility that participants optimize learning imprecisions and exploration in terms of a quantitative comparison between the costs (in terms of reward loss) and benefits (in terms of reduced cognitive resources for learning imprecisions, and of lower uncertainty for exploration). Together, these theoretical considerations suggest that the opposite adjustments of learning imprecisions and exploration to unexpected uncertainty follow the opposite changes in their reward costs between conditions.

## Discussion

Making variable decisions under uncertainty has clear benefits, but also significant costs (*9, 22*). Previous research has provided compelling evidence that part of this variability reflects active and adaptive exploration aimed at reducing uncertainty about choice options (*3*–*5*). However, seeking information about uncertain options often means foregoing a rewarding option (*10*). Furthermore, recent work has shown that a substantial fraction of decision variability arises not only from exploration, but also from learning imprecisions (*6*). Here, we studied how humans adapt these two sources of decision variability to different forms of uncertainty. We obtained converging evidence for an independent tuning of learning imprecisions and exploration to their specific costs and benefits. We discuss below the implications of these findings for existing theories of the explore-exploit trade-off that ignore learning imprecisions (*23*).

In agreement with existing theories (*4, 10, 14, 22*), we found that humans adjust the explore-exploit trade-off depending on the dominant form of uncertainty in their environment. Participants made more variable choices in the volatile condition where option values change over the course of a single game. Such ‘random’ exploration is adaptive in this context to monitor possible changes in the values of recently unchosen options (*3*). Similar adaptations of the explore-exploit trade-off have been described across other task conditions. Humans make little to no exploratory choices when the outcomes of unchosen options are known (*6*), engage in ‘directed’ exploration in conditions with imbalanced uncertainty across options (*5, 22, 24*), and use structured exploration in environments with spatially correlated option values (*25*).

Here, by measuring imprecisions in the reinforcement learning process that updates option values, we found that humans increase learning precision in the volatile condition. This within-participant adjustment of learning precision is resource-efficient: while outcomes from stochastic options are weakly informative about their values, outcomes from volatile options are informative about changes in their values. Participants therefore require not only higher learning rates, but also more precise updates. In the noisy Kalman filter that we fitted to participants’ behavior, the learning rate *α* reflects the perceived volatility of choice options. We leveraged the large individual differences in learning rate in the volatile condition to validate a second prediction of our resource-efficient account: that individuals with high learning rates (i.e., who perceive options are more volatile) should (on average) learn option values more precisely than individuals with low learning rates. This pervasive relation between learning rate and learning precision reveals a second trade-off that shapes human learning and decision-making under uncertainty.

Adjustments of exploration and learning imprecisions are not only distinct in terms of their respective costs and benefits. They also correspond to fundamentally different types of adjustments in the decision process. Optimizing the explore-exploit trade-off consists in tuning the choice policy between available options (*4*), downstream from the reinforcement learning process which estimates option values based on obtained rewards. By contrast, optimizing the cost-benefit trade-off associated with imprecise computations consists in tempering with the estimation of option values themselves, upstream from the choice policy (*6, 9*). The observation of opposite adjustments of exploration and learning imprecisions in response to unexpected uncertainty provides empirical evidence that humans can simultaneously and independently regulate how they choose between options, and how precisely they learn from choice outcomes.

Importantly, the tuning of learning imprecisions to uncertainty cannot be described in terms of ‘efficient coding’ theories (*7, 26*–*28*). These influential theories explain how noise-free computations can be tuned at long timescales to minimize the impact of external noise on performance. Instead, we show that humans rely on imprecise computations to learn option values and that they regulate this internal noise (*6,9*) at short timescales as a function of task demands. This finding is in agreement with the idea that humans optimize the use of their limited cognitive resources in a flexible, context-dependent fashion as a function of task demands (*29, 30*). The fact that humans regulate learning imprecisions confirm theoretical considerations that describe precise computations as extremely costly in terms of neural resources (*13*). This finding is also consistent with the observation that reinforcement learning in multidimensional environments relies on selective attention mechanisms that focus learning resources on a subset of learnable option features (*31*– *33*).

The fact that humans adapt their exploration and learning imprecisions across conditions does not imply that participants are aware and in control of these adaptations. In our computational model, learning imprecisions corrupt value representations in an implicit fashion – i.e., trial-to-trial learning errors are not explicitly accounted for by increasing the uncertainty about option values. It is well known that humans exhibit partial blindness to internal sources of error (*34, 35*). In additional control analyses (see *Supplementary Text*), we have considered variants of our learning agent where the uncertainty triggered by learning imprecisions alters learning rates. The fact that these variants provide a poorer fit to human decisions suggests that learning imprecisions may be regulated in an implicit, non-intentional fashion. By contrast, exploration is thought to reflect a source of decision variability that can be intentionally regulated – in particular in conditions where sources of uncertainty are known (as it is the case in our study). And in our model, exploration corresponds to explicit ‘non-greedy’ decisions that do not maximize expected value. Nevertheless, further work will be required to determine the extent to which exploration and learning imprecisions are regulated intentionally as a function of uncertainty.

We simulated the marginal reward costs of exploration and learning imprecisions in each condition, expressed in terms of relative loss compared to a learning agent without this source of variability. We found that unexpected uncertainty is associated with lower costs of exploration but higher costs of learning imprecisions than expected uncertainty. These opposite effects of unexpected uncertainty on the costs of exploration and learning imprecisions provide an adaptive account of their opposite adjustments. In agreement with this view, the marginal reward costs of participants’ exploration and learning imprecisions were found to be constant across conditions. It remains however unclear whether participants actively maintain fixed reward costs across conditions, or whether they optimize other cost-benefit trade-offs associated with these two sources of behavioral variability (see *Supplementary Text*).

If humans regulate (intentionally or not) the variability of their decisions in terms of two distinct cost-benefit trade-offs, then their neurophysiological substrates should be dissociable. Trial-to-trial decision variability has been associated with brain activity in frontopolar (*3, 37*) and anterior cingulate (*38, 39*) cortices, and related to dopaminergic and noradrenergic pathways (*40*–*43*). These multiple effects have been linked to adjustments of the explore-exploit trade-off, which is the sole source of decision variability in state-of-the-art models relying on perfectly precise computations. Accounting for learning imprecisions has revealed that the variability of value updates correlates with BOLD activity in the anterior cingulate cortex and with pupil dilation (*6*). Pupil dilation correlates with the activity of the locus coeruleus (*44, 45*), a brainstem nucleus with norepinephrine-containing neurons which has bidirectional projections with the anterior cingulate cortex in the primate brain (*46*). Adjustments of exploration and learning imprecisions could be achieved either by dissociable brain regions and neuromodulatory pathways, or by different signals in the same brain region (*47*). In particular, anterior cingulate activity has been related not only to exploration, but also to the cost-benefit trade-off associated with cognitive control (*12*). Future research should study how exploration and learning imprecisions are simultaneously regulated in the human brain (*48*).

The delineation of opposite adjustments of exploration and learning imprecisions in response to volatility suggests that models lacking either source may draw incorrect interpretations. Indeed, our findings show that humans adjust not only how much they explore uncertain choice options, by arbitrating the explore-exploit trade-off, but also how precisely they learn their values through reinforcement learning. These adjustments may generalize to different tasks and computational models of behavior. Indeed, decisions in perceptual categorization tasks suffer from a similar source of imprecisions in the underlying inference problem (*49*). These categorization tasks can be embedded in volatile conditions (*50*) where statistical inference models of behavior include a parameter that controls learning rate and another parameter that controls inferential imprecisions (*51, 52*). In this context that differs widely from the two-armed bandit used in this study, the same negative relation between learning rate and inferential imprecisions may emerge. Taken together, these results indicate that the precision of cognitive computations acts as an important and flexible constraint on human cognition (*9*). Existing theories of cognitive control should therefore be revised to account for imprecise computations and their associated costs and benefits.

## Materials and Methods

### Experimental design

Three samples of *N* = 200 English-speaking adults were recruited on Prolific (prolific.co) for inclusion in the two datasets collected for this study (a ‘discovery’ dataset and a ‘replication’ dataset, total *N* = 400). We fixed this sample size of *N* = 200 per dataset a priori and calculated that it is sufficiently powered to detect a minimal between-subject correlation of 0.20 with 80% power at a significance threshold *α* = 0.05. We excluded from analyses any participant whose accuracy did not significantly exceed chance-level performance at a one-tailed threshold *α* = 0.05 in any of the three conditions (binomial test against 50% accuracy). This exclusion procedure left *N* = 154 participants (mean age = 29.6 ± 10.0 years, 53 females) in the first dataset, and *N* = 142 participants (mean age = 25.4 ± 6.4 years, 72 females) in the second dataset. All participants provided informed consent regarding their participation in the study, which followed guidelines approved by the ethical review committee of the Institut National de la Santé et de la Recherche Médicale (IRB #00003888).

Upon gaining access to the task, participants were prompted to set the task to full-screen before being able to continue. Next, participants were introduced to the mechanics of their responses (e.g. keys for choosing options) and the presentation of the task as a slot machine game. They were instructed on the dynamics of the reference (Ref), stochastic (S+) and volatile (V+) conditions and the difference between these conditions via written prompts on the screen. Participants always played the Ref condition first, and the order of the other two conditions was counterbalanced across participants. Before every condition, participants played a ‘practice’ block to familiarize them with the condition. At the end of each practice block, participants were presented with a debriefing of their performance and the condition with a visual illustration of the trajectory of their decisions and the rewards they obtained. The task was self-paced but had to be completed within 60 minutes to avoid disengagement. Participants were offered to take breaks between blocks.

### Task generation

The Ref and S+ (stochastic) conditions corresponded to two-armed bandits with fixed reward distributions, drawn between 1 and 99 points. The V+ (volatile) condition corresponded to a restless two-armed bandit task where the means of the reward distributions associated with each option follow a random walk over time. In both conditions, the mean rewards 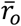 associated with the two options *0* ∈ {1,2} were symmetrical (i.e., 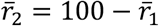). Reversals were defined as time points where the mean rewards associated with the two options cross each other.

Task difficulty was matched between the S+ and V+ conditions based on the simulated performance of an optimal learning agent. For each participant, we first generated 2,000 random walks of mean rewards following a beta distribution of initial mean of 50 points and a drifting standard deviation of 5 points. All random walks that travelled below 10 points or above 90 points, or had steps larger than 10 points, were discarded. Excursions below or above 50 points of 8, 12, 16, 20, and 24 steps were chosen to populate the V+ (volatile) condition. The average reward of each excursion was used as the static mean reward of a round of trials for the Ref (reference) condition. We set the variance of sampled rewards such that 75% of sampled rewards from the better option in the Ref condition were higher than 50 points (and 25% lower than 50 points). We then computed the effective sampling variance (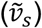) and the effective drifting variance (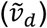) of rewards, and simulated a standard Kalman filter with greedy choices in the V+ condition to obtain its accuracy (fraction of choices toward the option associated with the largest mean reward). To generate the last, S+ (stochastic) condition, we incrementally regressed the static mean reward from the Ref condition toward 50 until the choice accuracy of a Kalman filter matched that of the V+ condition (Supplementary Figure 8). Participants played two instances of each condition (2 blocks of 80 trials for each condition).

Options were depicted as black shapes for the Ref and S+ conditions, and as colored discs in the V+ condition, to help participants distinguishing between stable (Ref and S+) and volatile (V+) conditions. To make sure that participants treated each new round of the Ref and V+ conditions as independent from the previous ones, we presented novel shapes for each new round. After each choice, the chosen option briefly appeared at the center of the screen, and was then followed by the number of points obtained from the chosen option (Figure 1A). Participants earned £3.30 upon successful completion of the task. Participants gained an extra £1.00 as a bonus if they performed significantly better than chance across the entire experiment (*α* = 0.05, meaning a choice accuracy above 53.75%). Participants who answered a battery of mental health questionnaires performed after the task (not analyzed for this study) earned another £3.00 upon completion.

The visuals and dynamics of the task (i.e., the frontend code) were implemented using jsPsych (version 6.3.0) (*53*). The backend server was handled using Node.js. Participants’ responses to the task and questionnaires were stored in a MySQL database.

### Computational modeling

Following each outcome, the posterior value of the chosen option 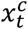 is updated according to a standard Kalman filtered corrupted by additive random noise *ε*_*t*_ drawn from a normal distribution of zero mean and standard deviation *η*_*t*_:

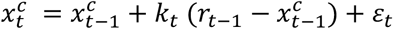

where *k*_*t*_ is the Kalman gain, which depends on the posterior variance of the chosen option 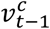 and the sampling variance *v*_*s*_ of rewards, which was set to its true effective value (0.0163 for rewards rescaled between 0 and 1) and used as ‘scaling parameter’ in the model:

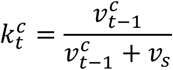

The posterior variances of the two options *0* ∈ {1,2} were updated using the standard equation of the Kalman filter:

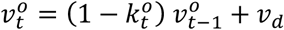

where *v*_*d*_ is the drifting variance assumed by the Kalman filter. We fitted *v*_*d*_ in terms of its associated asymptotic Kalman gain *α* for an option that would be chosen on *n* → ∞ trials. Following recent findings (*6*), the standard deviation of the random noise corrupting the update of the posterior value of the chosen option scales as a constant fraction *ζ* of the update 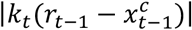. Posterior variances of the two options were initialized to the true variance of mean reward values across all conditions (0.0214 for rewards rescaled between 0 and 1).

The posterior value of the unchosen option 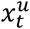 decays toward the baseline value (50 points, i.e., 0.5 for values rescaled between 0 and 1). The rate of this decay is controlled by an exponential decay factor *δ*:

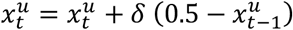

The probability of choosing option 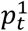 based on the posterior values of the two options follows the standard softmax policy with choice temperature *τ*:

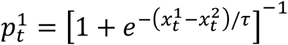

where *τ →* 0 corresponds to a purely greedy, argmax policy.

We used Sequential Monte Carlo (SMC) sampling methods to estimate the conditional likelihoods of the responses of each participant given the set of parameter values. Using Bayesian Adaptive Direct Search (BADS) with 10 random starting points for each parameter *α, ζ, δ* and *τ* (*54*), we first obtained point estimates of best-fitting parameter values. We then used these estimates as starting points for the estimation of their joint posterior distribution using Variational Bayes Monte Carlo (VBMC) (*55, 56*). Both model fitting algorithms accounted for noise in log-likelihood estimates due to the presence of random learning noise in the Kalman filter updates.

### Computational model validation

To ensure that the inclusion of learning noise (controlled by its Weber fraction *ζ*) and soft choices (controlled by their temperature *τ*) were both necessary to fit participants’ choices, we performed random-effects Bayesian Model Selection (BMS) based on its standard Dirichlet parameterization described in the literature (*57*). In practice, as in recent work (*6*), we compared the model including both sources of decision variability with two model variants: one without learning noise where *ζ* = 0, and one with a purely greedy choice policy where *τ* = 0. The full model outperformed the other two reduced models (model prevalence of 70% for the full model, exceedance *p* > 0.999).

Critically, we performed standard model recovery to validate our model simulation and fitting procedure (*20, 21*). In particular, by simulating the different variances of our Kalman filter model using individual best-fitting parameters, we established that our three candidate models were indeed distinguishable between each other (Supplementary Figure 2B). Our fitting procedure was not biased in that we were able to successfully recover all three model variances by simulating decisions from each model and then fitted all three models again to this simulated behavior for which the ground-truth model structure is known.

### Statistical analyses

We employed Wilcoxon signed-rank tests for the comparison of model-free behavioral variables across conditions (choice accuracy and switch rate). Wilcoxon signed-rank tests were also used for comparisons of best-fitting model parameters across conditions. Repeated-measures ANOVAs were performed for comparing choice accuracy curves and switch rate curves across conditions. Spearman (rank-based) correlations were used unless noted otherwise. Split-plot tests were used to calculate the effects of median splits with respect to each model parameter on choice accuracy and switch rate curves.

We performed a principal component analysis (PCA) on standardized (zero-mean and unit-variance) model parameter values such that, by construction: 1. the variance explained by a principal component (PC) in a given condition captures shared variance between model parameters, and 2. different PCs are orthogonal to each other and therefore reflect uncorrelated sources of variability in model parameters.

Standard errors on the Spearman’s rank correlation coefficients were generated through bootstrapping, by randomly sampled datasets from the original dataset (with identical sample size and replacement) 1,000 times and calculating the correlation on the new datasets. The same procedure was done to obtain the standard errors of PC coefficients and associated fractions of variance explained. Statistical significance of the fraction of variance explained was obtained by a one-sided comparison of the variance explained by bootstrapped PCs to the variance explained by bootstrapped PCs for shuffled data. A similar procedure was used to compare the fractions of variance of each PC explained by each model parameter.

## Acknowledgments

This work was supported by a starting grant from the European Research Council (ERC-StG759341) awarded to V.W., and by a department-wide grant from the Agence Nationale de la Recherche (ANR-17-EURE-0017, EUR FrontCog). M.R. is the beneficiary of a postdoctoral fellowship from the AXA Research Fund. M.R. is supported by La Fondation des Treilles. The funders had no role in study design, data collection and analysis, decision to publish or preparation of the manuscript. J.K.L. designed the experiments, conducted the experiments, analyzed the data and wrote the paper. M.R. provided feedback on data analyses and wrote the paper. V.W. designed the experiments, analyzed the data, wrote the paper and supervised the study. The authors declare that they have no competing interests. The datasets generated and analyzed during the study will be made freely available on an online repository. The analysis code supporting the reported findings is currently available from the corresponding authors upon request and will be made freely available on an online repository upon publication of the manuscript.

