## Supplementary Materials for "Adaptive tuning of human learning and choice variability to unexpected uncertainty"

**This PDF file includes:**

Supplementary Text (pages 1-6)

Supplementary Figures 1 to 9 (pages 7-15)

#### Supplementary Text

##### Comparison between Kalman filter and Reinforcement Learning (RL) models

In this study, we used a Kalman filter (Kalman, 1960) to track the statistics of option values for two main reasons:

1. Unlike simpler reinforcement learning (RL) models, the instantaneous learning rate of the Kalman filter (referred to as its Kalman gain) varies from trial to trial as a function of the estimated uncertainty regarding the value of the chosen option. This allows the Kalman filter to have decreasing instantaneous learning rates as the statistics of options with constant mean values (i.e., no unexpected uncertainty) are learnt. Note that this is not the case for the simpler reinforcement learning (RL) models based on the Rescorla-Wagner rule, which use fixed learning rates over time (*6, 58, 59*).

   The constant learning rate used by canonical RL models fitted to human data is particularly problematic in the Ref and S+ conditions, where participants are instructed explicitly that the two options are associated with constant mean values (i.e., no unexpected uncertainty) - and where learning rates should decrease over time in each new round of trials.

   By contrast, by monitoring not only the means of option values but also their variances, a Kalman filter can accurately track option values whose means can either remain constant (as in the Ref and S+ conditions) or vary (as in the V+ condition) across successive trials. This theoretical advantage motivated the use of a Kalman filter to fit human choice data in our task. Note that the update equation of the Kalman filter is highly similar to the Rescorla-Wagner rule, except that its instantaneous learning rate varies from trial to trial.

We have fitted the noisy RL model described in previous work to human data (*6*) and found that the Kalman filter provided a better fit of human choices in all three conditions. We adopted here a fixed-effects analysis, by summing the log-marginal likelihood of participants to assess whether the group of participants as a whole was better fitted by the noisy Kalman filter ($H_1$) or by the noisy RL model ($H_0$). We obtained decisive evidence in favor of the noisy Kalman filter in each condition (Ref: $\log(BF_{10}) = +95.2$; S+: $\log(BF_{10}) = +91.2$; V+: $\log(BF_{10}) = +38.9$).

2. The interpretability of the asymptotic learning rate of the Kalman filter is much higher than the constant learning rate of the canonical RL model. Indeed, the asymptotic learning rate $\alpha$ of the Kalman filter is a monotonically growing function of the structure of uncertainty assumed by the learning agent:

   $$\alpha = F(v[\text{drift}]/v[\text{sample}])$$

   where $v$[drift] is the variance of the drift in option values across successive trials (i.e., unexpected uncertainty), whereas $v$[sample] is the variance of reward samples around their corresponding option value (i.e., expected uncertainty). F(x) is a sigmoidal function which can be accurately approximated as follows (see Supplementary Figure 9):

   $$F(x) = 1/(1+\exp(+0.4486-\log_2(x)*0.6282))\text{^}0.5057$$

   This means that we can compute normative values of $\alpha$ in each condition from the generative statistics of option values used in each condition. Again, this is not possible for simpler RL models. In practice, because option values have constant means in the Ref and S+ conditions, the normative $\alpha$ is exactly zero. In the V+ condition where option values have drifting means, the normative $\alpha$ is 0.668. We can then compare these normative values (for a theoretical learning agent without any of the cognitive limitations that we have implemented in the noisy Kalman filter) to the best-fitting values obtained from the human data.

**Alternative account of the relationship between learning rate and learning noise**

Our implementation of the noisy Kalman filter is blind to the presence of learning noise in its trial-to-trial adjustment of learning rate (i.e., the Kalman gain). We grounded this implicit assumption on recent work showing that while humans account optimally for sensory noise during decision-making, they are essentially blind to noise introduced by their own cognitive system (*35*).

There is an alternative implementation of the noisy Kalman filter that would account for the presence of learning noise in its trial-to-trial adjustment of learning rate. Therefore, as a control, we have implemented two distinct versions of the noisy Kalman filter which accounts for the presence of learning noise in its update equations.

In the first version, the presence of learning noise is accounted for in the update equation for the posterior variance $v[t]$, as follows:

$$v[t] = (1- k[t])\ v[t\text{-}1] + v[\text{drift}] + \sigma[t]^2$$

where $k[t]$ is the Kalman gain at trial $t$ and $\sigma[t]^2$ is the variance of the learning noise corrupting the update of the posterior mean at trial $t$. This means that the agent considers learning noise as increasing uncertainty about the posterior mean, and increases the associated posterior variance accordingly. In practice, this first version results in an increased Kalman gain on trial $t$+1 when $\sigma[t]$ is large – to account for the increased posterior variance $v[t]$. We fitted this alternative version of the noisy Kalman filter to the same data, and compared its associated log-marginal likelihood to the one obtained for the 'noise-blind' noisy Kalman filter reported in the main text. We applied a 'fixed-effects' approach to compare the hypothesis that all participants are drawn from the 'noise-blind' version of the model to the hypothesis that all participants are drawn from the model accounting for learning noise in the update equation for the posterior variance. We obtained decisive evidence in favor of the 'noise-blind' model reported in the main text, in all three conditions (Ref: log(BF) = +244.9; S+: log(BF) = +268.0; V+: log(BF) = +73.0).

This first version of the model that accounts for the presence of learning noise is grounded on the effective impact of learning noise on uncertainty about the current posterior mean (i.e., its associated variance), but in the case of Weber noise – whose standard deviation scales with the magnitude of updates – this implementation may generate a 'runaway loop' whereby the agent increases its Kalman gain on trial $t$+1 to mitigate the impact of learning noise on trial $t$, but ends up increasing the learning noise on trial $t$+1 (which scales with Kalman gain). We therefore considered a second version, which accounts for large learning noise by actively reducing its Kalman gain on the following trial.

In this second version, the presence of learning noise is accounted for in the update equation for the Kalman gain $k[t]$ itself, as follows:

$$k[t] = v[t\text{-}1] \,/\, (v[t\text{-}1] + v[\text{sample}] + \sigma[t\text{-}1]^2)$$

where $v[t\text{-}1]$ is the posterior variance at trial $t$-1 and $\sigma[t\text{-}1]^2$ is the variance of the learning noise corrupting the update of the posterior mean at trial $t$-1. This means that the agent actively reduces its Kalman gain following trials associated with large learning noise. Again, we fitted this alternative version of the noisy Kalman filter to the same data, and compared its associated log-marginal likelihood to the original model using the same 'fixed-effects' approach. We obtained clear evidence in favor of the 'noise-blind' model over this alternative implementation which accounts for learning noise in its Kalman gain equation (Ref: log(BF) = +4.3; S+: log(BF) = +14.3; V+: log(BF) = +110.3).

Importantly, the best-fitting parameter values of learning rate $\alpha$ and learning noise $\zeta$ showed the same differences between conditions in this alternative implementation of the noisy Kalman filter. In particular, the learning rate $\alpha$ was still substantially larger in the V+ condition than in the S+ condition (signed-rank test, $z = +9.8$, $p < 0.001$). And the learning noise $\zeta$ was still significantly lower in the V+ condition than in the S+ condition (signed-rank test, $z = -3.9$, $p < 0.001$). Finally, learning noise $\zeta$ still correlated negatively with learning rate $\alpha$ in each condition in this version of the model which accounts for the presence of learning noise in the update equation for the Kalman gain (rank correlation, Ref: Spearman's $\rho = -0.263$, $p < 0.001$; S+: Spearman's $\rho = -0.296$, $p < 0.001$; V+: Spearman's $\rho = -0.332$, $p < 0.001$).

Together, these results argue against the alternative interpretation that it is variations in learning noise across conditions and participants that trigger variations in learning rate, rather than the converse as we propose.

**Decay parameter**

The decay parameter $\delta$ can be considered either as a normative feature of the model in environments with mean-reverting statistics, or as a cognitive feature of the model corresponding to a decay of unchosen option values in working memory. Owing to the different conditions of our task, it is possible to arbitrate between these two views of the decay parameter $\delta$:

1. The decay parameter $\delta$ can be seen as a normative feature of the update equations for options whose values not only drift randomly but also regress toward a long-term mean (in this case, 0.5). Note, however, that this can only be the case in environments (conditions) with unexpected uncertainty. In our study, the Ref and S+ conditions are described explicitly to participants as conditions where option values are <u>constant</u> over time. If the decay parameter $\delta$ reflects participants' assumption of mean-reverting option values, then it should only be present (i.e., $\delta > 0$) in the V+ condition where option values are changing over time. In this case, the decay should theoretically be applied to both the chosen and the unchosen option values in the update equations of the noisy Kalman filter.

2. By contrast, the decay parameter $\delta$ can also be seen as a cognitive feature of the model, corresponding to a decay of unchosen option values in working memory toward a 'baseline' value (in this case, 0.5 = their overall mean). Under this second hypothesis, which is the one we have favored in our implementation of the noisy Kalman filter, the decay should be present (i.e., $\delta > 0$) not only in the V+ condition but also in the Ref and S+ conditions. In this case, the decay should only be applied to the unchosen option value in the update equations of the noisy Kalman filter.

A useful feature of our implementation of the decay is that the Kalman filter without decay is nested as a special case of the Kalman filter with decay (i.e., where $\delta = 0$). To test whether a decay is needed not only in the V+ condition where option values are changing over time, but also in the Ref and S+ conditions where option values are constant over time, we compared the version of the Kalman filter where $\delta = 0$ to the version (reported in the main text) where $\delta$ is fitted as a free parameter. In agreement with the idea that $\delta$ reflects a decay of unchosen option values in working memory rather than the assumption of mean-reverting option values, we found that the noisy Kalman filter with $\delta > 0$ provides a better fit of participants' choices in all three conditions. As above, we adopted a fixed-effects, and obtained decisive evidence in favor of the noisy Kalman filter with decay in all conditions (Ref: log(BF) = +107.3; S+: log(BF) = +107.9; V+: log(BF) = +716.1).

Note that a similar decay of unchosen option values has previously been proposed in the literature (*60*). These additional results clarify the nature of the decay parameter $\delta$ in the model, indicating that it reflects a suboptimal decay of unchosen option values in working memory rather than a normative adaptation of learning to mean-reverting option values under unexpected uncertainty.

**Theoretical efficiency of parameter adjustments between conditions**

We considered the use of relative reward loss (defined as a fraction of the reward excess that would have been obtained in the absence of learning noise or choice temperature) as a metric to measure 'efficiency.' Here, we explain how the decrease in learning noise $\zeta$ and the increase in choice temperature $\tau$ in the volatile condition are 'efficient'. We have performed additional analyses, indicating that these adjustments are not efficient in an absolute sense, when efficiency is based on the optimization of a cost-benefit trade-off that considers rewards (and their loss), but rather in a relative sense.

First, the idea that participants rely on a relative (condition-specific) estimation of reward loss – rather than the absolute reward loss – stems from a rich literature in human psychology. To cite only some of the most relevant literature for our study, value-based decisions are thought to be based on context-dependent representations of option values (*61, 62*). In our task, participants were cued at the beginning of each block about the upcoming condition, and they were instructed that the stochastic (S+) and volatile (V+) conditions were more difficult than the reference (Ref) condition. Participants could therefore estimate their performance in a condition-specific fashion.

The impact of learning noise and choice temperature in terms of absolute reward excess is shown in Supplementary Figure 7A (in contrast to relative reward excess as shown in Figure 7A). Plotting the derivatives of the reward loss curves defined in a relative sense as a fraction of the reward excess that would have been obtained in the absence of learning noise or choice temperature, separately for each condition, is shown in Supplementary Figure 7B.

Consider another specification of efficiency, as a cost-benefit optimization between the gain of having greater imprecisions (e.g., reduction in cognitive resources used for learning) against their costs (i.e., decreasing reward). Note that under this assumption, we measure the gain on the same scale as reward. Assume that this gain function is positive with increasing learning noise and has a monotonically decreasing derivative on the same domain. One can find such a function which can fit the three values of learning noise observed in the three conditions only when considering reward loss in a relative, condition-specific sense. In other words, our data are compatible with the idea that participants adjust learning noise to maximize efficiency where reward loss is defined in a relative, condition-specific sense. Because efficiency may be defined a priori in terms of absolute (not relative) reward, these adjustments of learning noise and choice temperature across conditions as adaptive but not necessarily as efficient. We also do not make strong claims regarding the specific arbitration that participants rely on to adjust these parameters. What we show unambiguously, is that volatility increases the relative cost of learning noise and decreases the relative cost of choice temperature.

The relative reward loss curves shown on Figure 7A show that volatility increases the relative cost of learning noise and decreases the relative cost of choice temperature: 1. any given level of learning noise has a higher relative cost in the volatile condition, and 2. any given level of choice temperature has a lower relative cost in the volatile condition. By adopting lower learning noise and higher choice temperature in the volatile condition, participants effectively achieve an approximately constant level of relative reward loss across conditions, which we show in Figure 7B. This observation is not constrained by theory, it is an observation derived from the data through simulations using the same levels of learning noise and choice temperature as participants. Conversely, adopting a constant level of relative reward loss across conditions results in lower learning noise and higher choice temperature in the volatile condition. These different observations, all dependent on data and simulations rather than on a strict definition of efficiency, all point toward the idea that the adjustments of learning noise and choice temperature are adaptive when considering reward loss in a relative sense.

The two-dimensional plots of Supplementary Figure 7C show that the volatile condition is associated with a larger partial derivative of the relative reward loss function $L^{rel}(\zeta, \tau)$ with respect to $\zeta$ for any value of $\tau$. , and a lower partial derivative of the same loss function with respect to $\tau$ for any value of $\zeta$. Another interesting observation is that participants achieve a constant level of relative reward loss $L^{rel}(\zeta, \tau)$ across all three conditions. This means that this consistency of relative reward loss across conditions holds whether we consider the reward loss triggered by a single parameter, or the reward loss triggered jointly by both parameters.

##### Relation between model-based PCA scores and model-free behavior

We conducted multivariate regressions of accuracy and switch rate in each condition as a function of standardized PC1 and PC2 scores. This analysis confirmed that the two PCA components have qualitatively distinct relations to behavior.

PC1 has a strongly negative relation to accuracy in the stable conditions (Ref: -0.051 ± 0.005, $t(151) = -10.7$, $p < 0.001$; S+: -0.041 ± 0.006, $t(151) = -6.8$, $p < 0.001$). This is highly expected because PC1 is associated with positive coefficients for learning rate and choice temperature, both of which are maladaptive when option values do not change over the course of a block. Interestingly, PC1 has a much weaker negative relation to accuracy in the volatile condition (V+: -0.019 ± 0.006, $t(151) = -3.2$, $p = 0.001$), which makes sense because high learning rates and choice temperatures can facilitate the learning of changing option values. PC1 has a strongly positive relation to switch rate in all three conditions (Ref: 0.073 ± 0.004, $t(151) = 20.1$, $p < 0.001$; S+: 0.086 ± 0.004, $t(151) = 23.5$, $p < 0.001$; V+: 0.068 ± 0.005, $t(151) = 14.5$, $p < 0.001$). This is expected because high learning rates and choice temperatures both contribute to switches between options.

PC2 has a milder negative relation to accuracy than PC1 in all three conditions (Ref: -0.026 ± 0.005; S+: -0.004 ± 0.006; V+: -0.018 ± 0.006). This is because PC2 reflects a trade-off between learning rate and learning noise: participants with high PC2 scores have higher learning rates (which is detrimental to accuracy) but lower learning noise (which is beneficial to accuracy). PC2 is therefore not strictly maladaptive in any conditions, unlike PC1 in the stable conditions. And unlike PC1, PC2 does not have a significant relation to switch rate in any condition (Ref: 0.004 ± 0.004, $t(151) = 1.0$, $p = 0.306$; S+: 0.004 ± 0.004, $t(151) = 0.9$, $p = 0.356$; V+: -0.003 ± 0.005, $t(151) = -0.7$, $p = 0.509$).

### Supplementary Figures

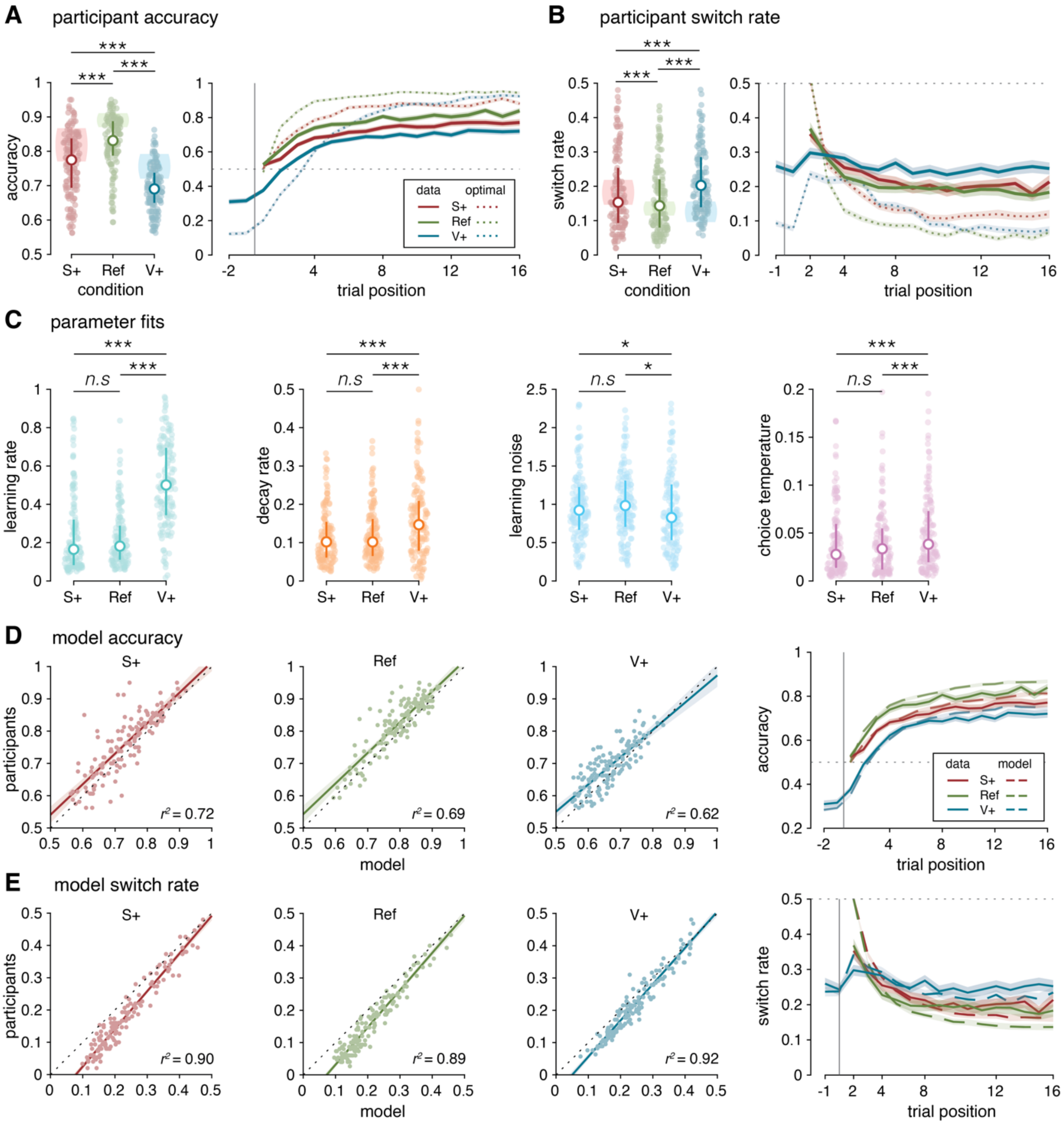


**Supplementary Figure 1. Behavioral and modeling results in the replication dataset**
**(A)** Average accuracy within each condition. Colored dots represent individual participants' mean accuracy. White dots indicate the median accuracy. Error bars represent the 1st and 3rd quartiles. Shaded areas indicate 1st and 3rd quartiles of the optimal model's accuracy. **(B)** Accuracy over time within each condition. Solid lines indicate the mean accuracy across participants. Dotted lines indicate the mean accuracy of the optimal model. The vertical line represents the start of a new block or a reversal. **(C)** Colored dots represent individual participants' fits for each parameter. White dots indicate the median fitted parameter value. Error bars represent the 1st and 3rd quartiles. **(D)** Left: Scatter plots of the average accuracy of participants to their best fitting models in each condition. Right: Accuracy over time within each condition. Solid lines indicate the mean accuracy across participants. Dashed lines indicate the mean accuracy of the best fitting model. **(E)** Left: Scatter plots of the switch rate of participants to their best fitting models in each condition. Right: Proportion of switches over time within each condition. Solid lines indicate the mean switch rate across participants. Dotted lines indicate the mean accuracy of the best fitting model. Shaded areas correspond to the SEM.

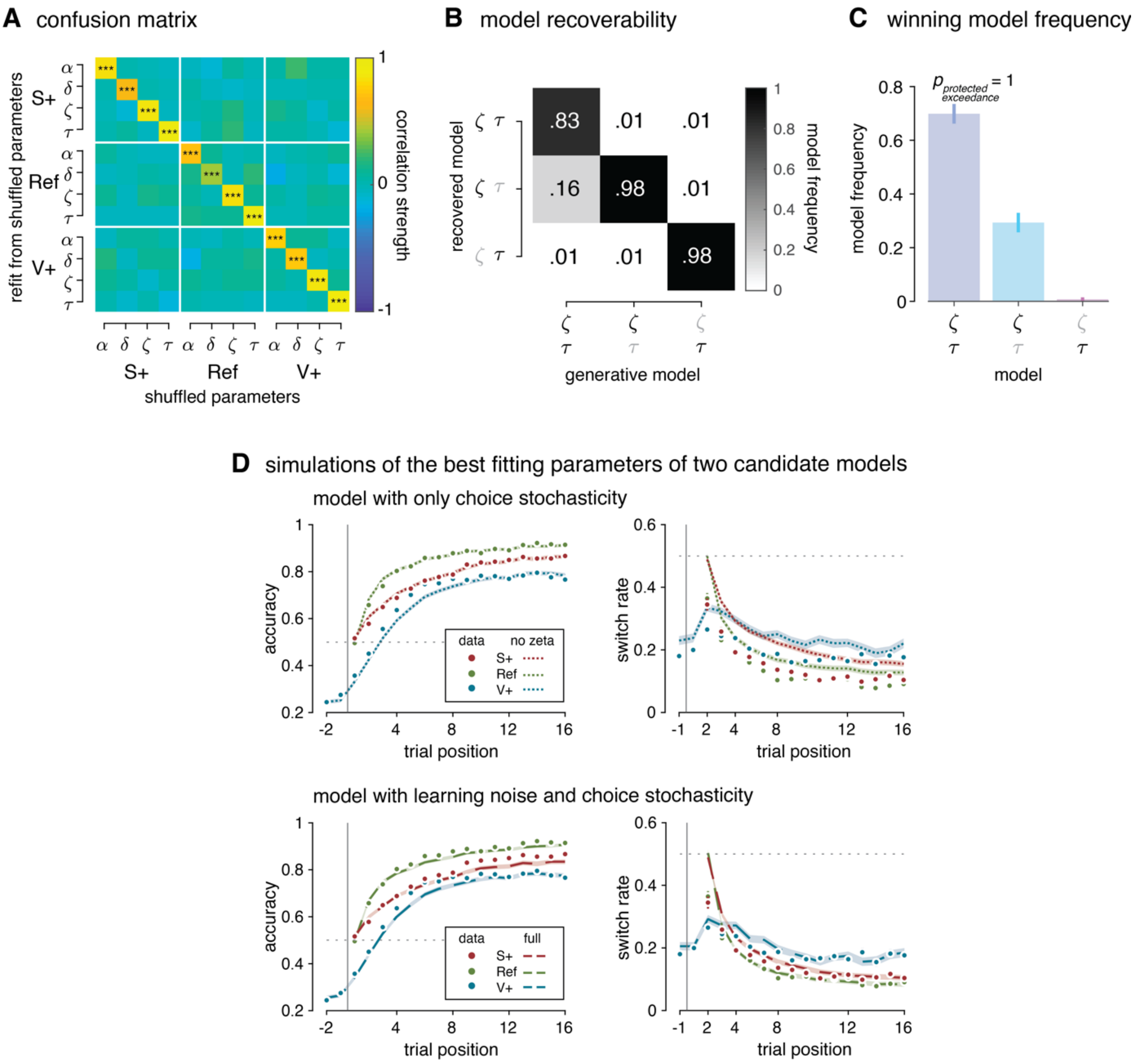


**Supplementary Figure 2. Computational model validation**
**(A)** Correlation matrix of parameters after fitting the model on data generated from the shuffled parameter set. **(B)** Frequencies of recovered model on simulated data from the three candidate models. **(C)** Winning model frequencies on participant data. **(D)** Accuracy and switch rate behavior from simulating two candidate models (KF model with and without learning noise, plus softmax) with their best-fitting parameters.

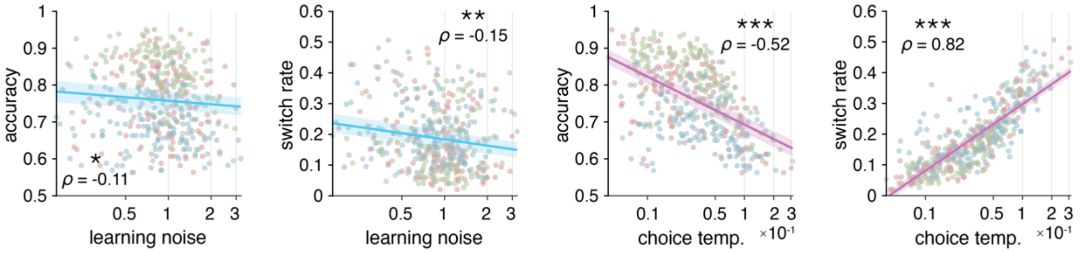


**Supplementary Figure 3. Behavioral expressions of learning and decision noise in the replication dataset**
Correlations of behavioral measures with learning noise and choice temperature in the replication dataset. Learning noise and choice temperature axes are spaced with log-scaling. Shaded areas are 95% CI.

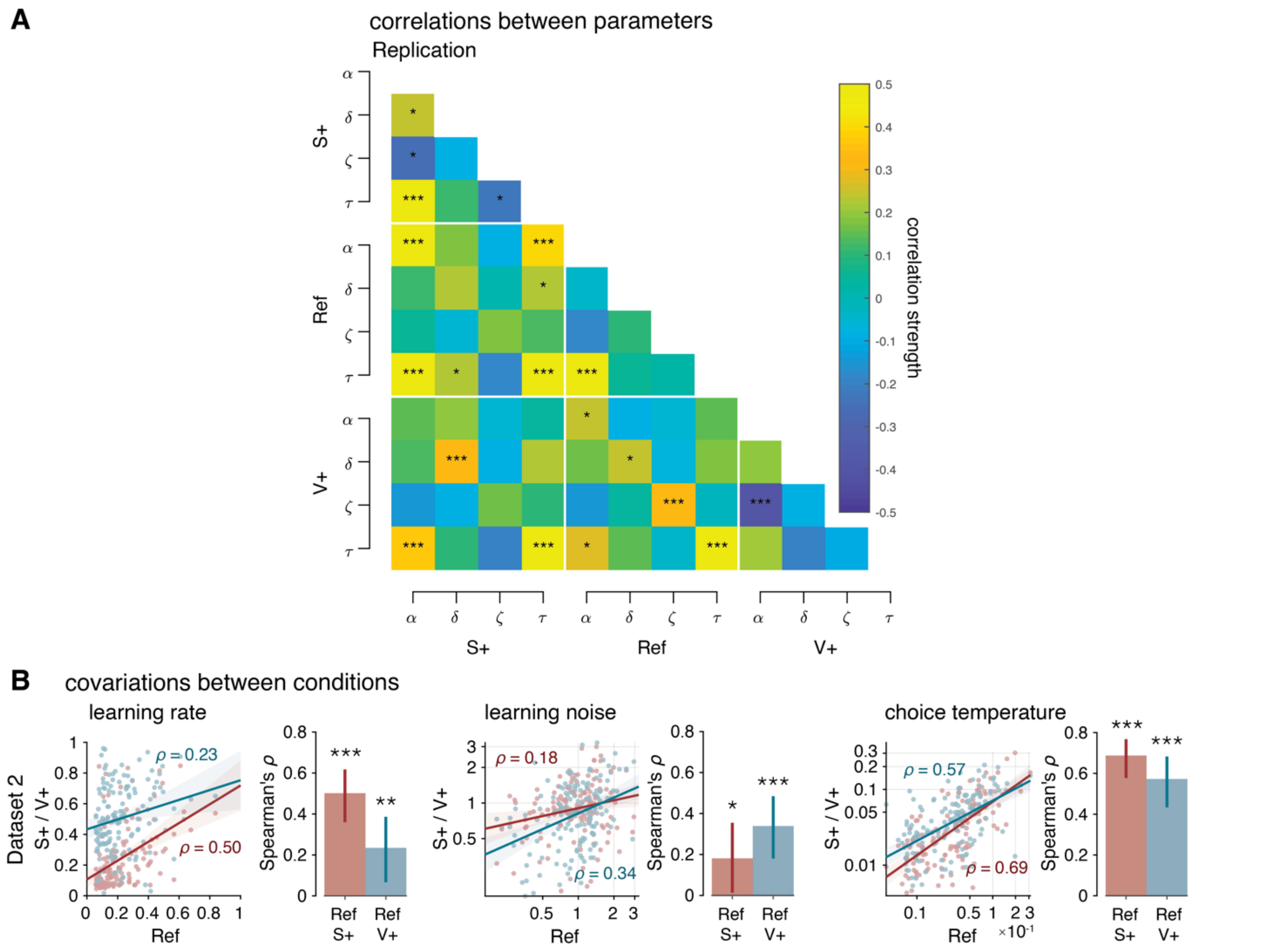


**Supplementary Figure 4. Parameter correlation matrices and covariations between conditions in the replication dataset**

**(A)** Left: Correlation matrix of participants' fitted parameter in the second dataset. Right: Correlation matrix of participants' fitted parameter in the third dataset. **(B)** Covariations of the learning rate, learning noise, and choice temperature between the reference (Ref) to the stochastic (S+) and the volatile (V+) conditions in the second and third datasets. Lines in scatter plots indicate the best-fitting regression line (blue: V+ to Ref; red: S+ to Ref). Shaded areas are 95% CI. Error bars on bar plots indicate 95% bootstrapped CI on the Spearman's ρ value.

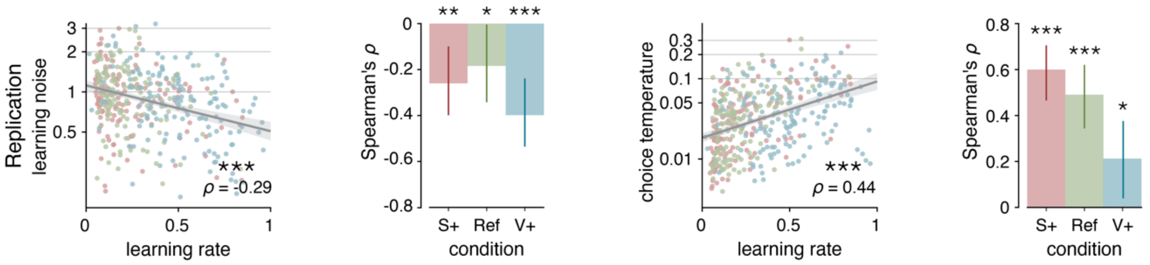


**Supplementary Figure 5. Covariations between parameters in the replication dataset**
**(A)** Correlations of learning noise and choice temperature with learning rate in the second and third datasets. Regression lines fitted on aggregate data pooling all conditions. Shaded areas are 95% CI. Error bars on bar plots indicate 95% bootstrapped CI on the Spearman's ρ value.

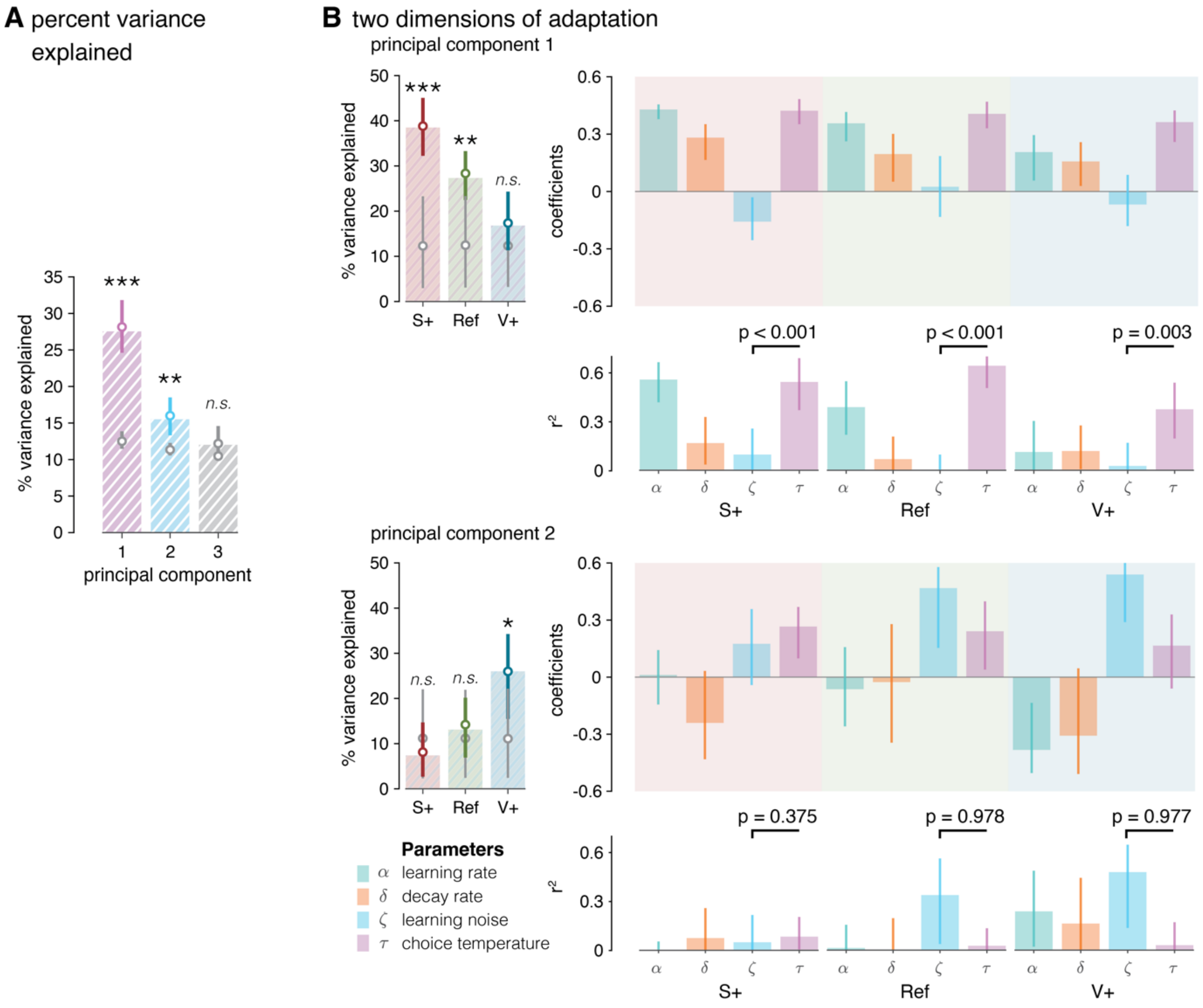


**Supplementary Figure 6. Principal component analysis on model parameters in the replication dataset**
**(A)** Percent variance explained up to the first non-contributive PC. Colored dots correspond to the median value of the percent variance explained from the bootstrap procedure. Gray dots are median values of the percent variance explained from PCs of shuffled data. Statistical significance calculated from one-tailed bootstrap significance tests. **(B)** Left: Percent variance explained by the first two PCs within each condition. The first PC, dominated by the variation of choice temperature, explains best the parameter adaptations in the non-volatile conditions. The second PC, dominated by the variation of learning noise, explains best the parameter adaptations in the volatile condition. Upper right: Ingredients and coefficients of the first two principal components. Lower right: Coefficient of determination of each parameter for the PC. All error bars are 95% bootstrapped CI.

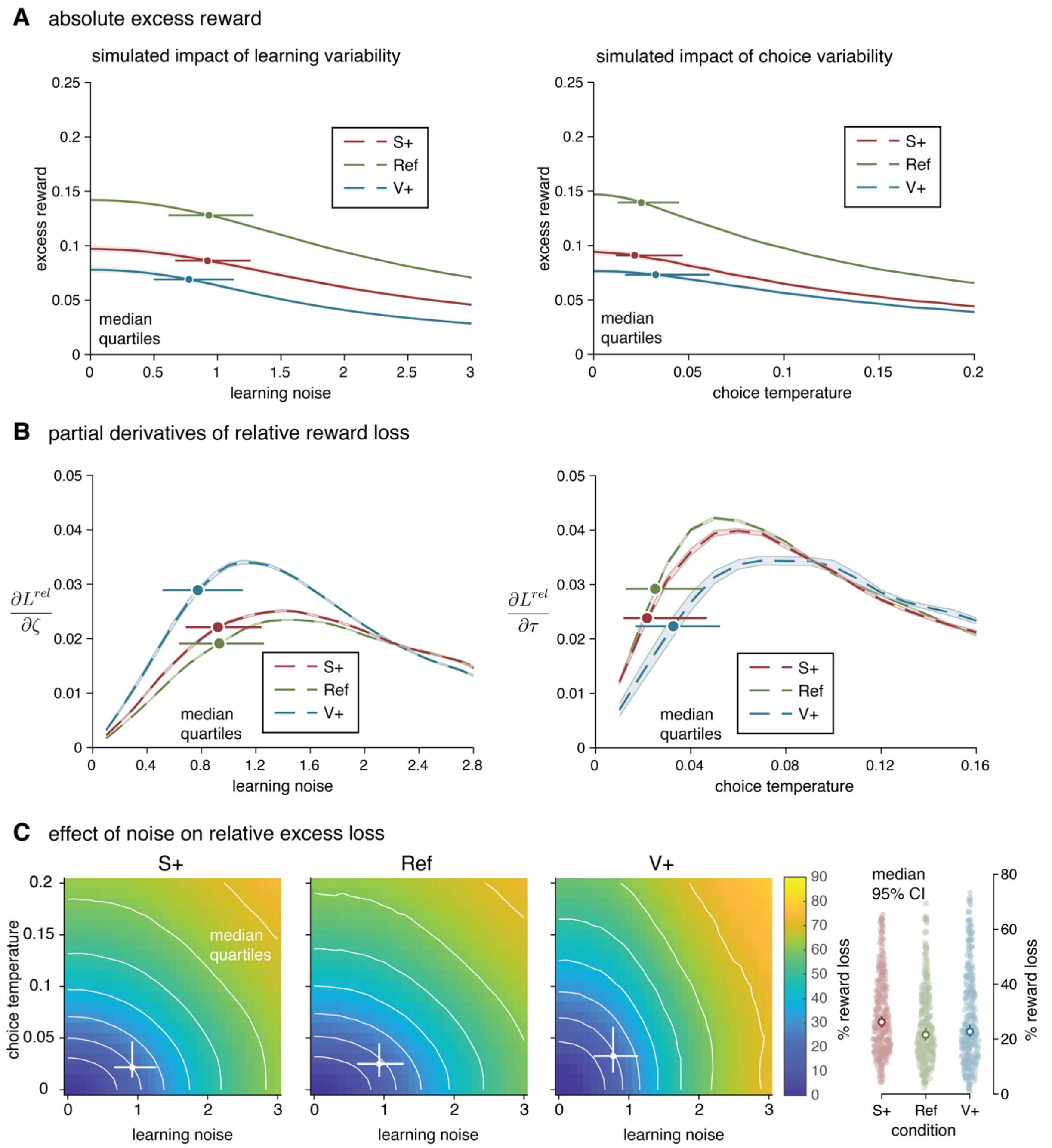


**Supplementary Figure 7. Absolute and relative reward loss simulations**
(**A**) Simulated absolute marginal rewards costs associated with each source of decision variability through simulations of the suboptimal learning agent by varying selectively the associated model parameter. (**B**) Partial derivatives of the relative reward loss from Figure 7A. Circles indicate median values of participants' parameter fits. Lines indicate 1$^{st}$ and 3$^{rd}$ quartiles. (**C**) Joint distribution of the relative reward costs associated with both sources of decision variability. Circles indicate median values of participants' parameter fits. Lines indicate 1$^{st}$ and 3$^{rd}$ quartiles.

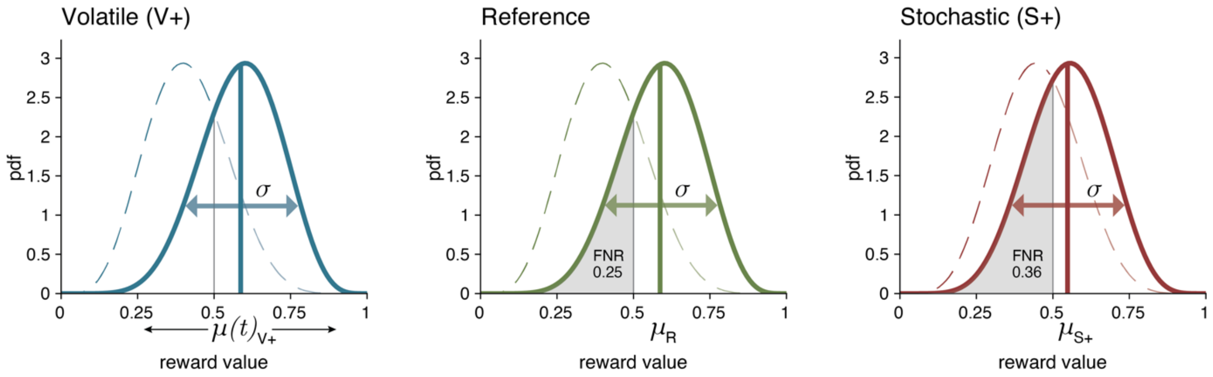


**Supplementary Figure 8. Generative distributions of the reward in each condition**
For each instance of the task, the reward of the best option in the Reference condition from a Beta distribution with shape parameters α, β, such that the mean and variance of the distribution are $\mu_R$ and σ with the false negative rate (FNR) of positive rewards being 25%. The reward of an option in the Volatile condition is generated by sampling at each trial from Beta distributions with shape parameters that correspond to a mean following a random walk, $\mu_{V+}(t)$, and variance σ as before. The reward for the best option in the Stochastic condition is generated by regressing the static mean, $\mu_R$, toward 0.5 to obtain $\mu_S$, such that the accuracy from the reward trajectories in V+ matches that of S+. The reward scale in all distributions is scaled down by a factor of $10^2$ (rewards seen in task are from 0 to 100).

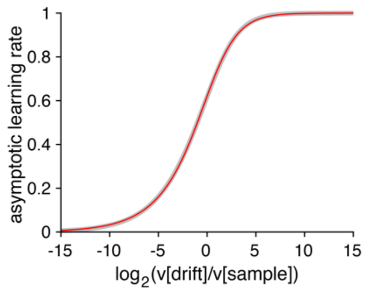


**Supplementary Figure 9. Relation between the structure of uncertainty assumed by the learning agent and the asymptotic learning rate of the Kalman filter.** The structure of uncertainty is expressed as $\log_2(v[\text{drift}]/v[\text{sample}])$. The gray line corresponds to numerical simulations of Kalman filters after $N$ = 1,000 observations, whereas the red line corresponds to the functional approximation described above.
